# Comprehensive scanning mutagenesis of human retrotransposon LINE-1 identifies motifs essential for function

**DOI:** 10.1101/721357

**Authors:** Emily M. Adney, Matthias T. Ochmann, Srinjoy Sil, David M. Truong, Paolo Mita, Xuya Wang, David J. Kahler, David Fenyö, Liam J. Holt, Jef D. Boeke

**Affiliations:** Institute for Systems Genetics and Department of Biochemistry Molecular Pharmacology, NYU Langone Health, New York, NY 10016; McKusick-Nathans Institute of Genetic Medicine, Johns Hopkins University School of Medicine, Baltimore, MD 21205; Division of Medical Biotechnology, Paul Ehrlich Institute, 63225 Langen, Germany; High Throughput Biology Laboratory, NYU Langone Health, New York, NY 10016

**Keywords:** LINE-1, L1, retrotransposon, scanning mutagenesis

## Abstract

Long Interspersed Nuclear Element-1 (LINE-1, L1) is the only autonomous active transposable element in the human genome. The L1-encoded proteins ORF1p and ORF2p enable the element to jump from one locus to another via a “copy and paste” mechanism. ORF1p is an RNA-binding protein and ORF2p has endonuclease and reverse transcriptase activities. The huge number of truncated L1 remnants in the human genome suggests that the host has likely evolved mechanisms to prevent full L1 replication and thereby decrease the proliferation of active elements and reduce the mutagenic potential of L1. In turn, L1 appears to have a minimized length to increase the probability of successful full-length replication. This streamlining would be expected to lead to high information density. Here, we describe the construction and initial characterization of a library of 538 consecutive trialanine substitutions that scan along ORF1p and ORF2p to identify functionally important regions. In accordance with the streamlining hypothesis, retrotransposition was overall very sensitive to mutations in ORF1p and ORF2p, only 16% of trialanine mutants retained near-wild-type activity. All ORF1p mutants formed near-wild-type levels of mRNA transcripts and seventy-five percent formed near-wild-type levels of protein. Two ORF1p mutants present a unique nucleolar-relocalization phenotype. Regions of ORF2p that are sensitive to mutagenesis, but lack phylogenetic conservation were also identified. We provide comprehensive information on the regions most critical to retrotransposition. This resource will guide future studies of intermolecular interactions that form with RNA, proteins and target DNA throughout the L1 life cycle.

## Introduction

Approximately 45% of the human genome consists of retroelements, three of which are highly active non-LTR retrotransposon families in modern humans: L1, Alu and SVA. These mobile genetic elements use a “copy and paste” mechanism called retrotransposition to propagate themselves within the host genome. The long interspersed element-1s (LINE-1s or L1s) are the only autonomously active human mobile element (Ostertag and Kazazian 2001; Brouha et al. 2003). Alu and SVA elements depend on L1-encoded proteins to execute retrotransposition and are thus considered non-autonomous.

There are roughly 500,000 copies of L1, making up about 17% of the human genome (Lander et al. 2001). The vast majority of these are severely 5’ truncated, and have diverged from the L1 consensus sequence, suggesting that they are very old and incapable of retrotransposition (Szak et al. 2002; Beck et al. 2010). About 15% of genomic L1Ta copies are full-length (Szak et al. 2002) and 6% of newly recovered experimentally induced elements were full-length (Symer et al 2005, Gilbert et al 2005, Gilbert et al 2007), but the latter value is probably an undercount due to less efficient recovery of full-length elements. Nevertheless, approximately 90 L1 elements per diploid human genome remain retrotransposition-competent and ongoing L1 activity continues to shape the evolution of mammalian genomes (Kazazian 2004; Huang et al. 2012; Faulkner and Garcia-Perez 2017).

The enormous number of 5’ truncated LINEs is a genomic feature of diverse species but despite this, is not well understood mechanistically. The pervasiveness of 5’ truncation may reflect the action of anti-retrotransposon factors that play an active role in minimizing retrotransposon length. If these assumptions are correct, minimization of L1 length might help reduce the opportunity for truncations. As a consequence, L1 would become streamlined and highly enriched for sequences key for retrotransposition.

L1 activity plays important roles in both normal development and pathology. There is evidence that L1 activity is highest in the germline and somatic insertion events are also reported in a variety of tissues, notably the brain, as well as during early development (Ostertag et al. 2002; Muotri et al. 2005; An et al. 2006; Kano et al. 2009; O’Donnell et al. 2013; Carreira et al. 2014). Insertions into coding regions can cause human disease (Hancks and Kazazian 2016) and increased L1 expression (and in some cases retrotransposition) is also observed in various cancers (Lee et al. 2012; Rodic et al. 2014; Doucet-O’Hare et al. 2015; Ardeljan et al. 2017; Burns 2017; Nguyen et al. 2018) L1 activity has been reported to correlate with aging, stress, DNA damage, and telomere shortening, all of which are processes that are likely normally regulated to keep the mutagenic capacity of L1 jumping in check (Gorbunova et al. 2014; Van Meter et al. 2014; De Cecco et al. 2019). Therefore, understanding of the mechanisms of L1 retrotransposition provides insight and opportunities in the fields of genome evolution, development, cancer biology, aging and neurodegeneration.

The full-length human L1 element specifies production of a 6 kb long transcript that encodes two proteins, ORF1p and ORF2p (Ostertag and Kazazian 2001), which are both essential for retrotransposition. ORF1p is a 40 kDa protein with both nucleic acid-binding and chaperone activities (Kolosha and Martin 1997; Martin and Bushman 2001). ORF2p is a 150 kDa protein that has endonuclease (Feng et al. 1996), reverse transcriptase (Mathias et al. 1991), and nucleic acid binding (Piskareva et al. 2013) activities. Upon translation of L1, ORF1p and ORF2p are thought to bind the same RNA molecule from which they were transcribed through a poorly understood process called cis-preference, also thought to require the 3’ poly(A) tail of L1 RNA (Boeke 1997; Wei et al. 2001; Kulpa and Moran 2006; Doucet et al. 2015). ORF1p is translated quite efficiently, but ORF2p translation occurs at much lower levels, through an unconventional process that is also poorly understood (Alisch et al. 2006). The L1 RNA, ORF1p, ORF2p, complex is referred to as the L1 ribonucleoprotein (RNP) complex and is likely to be the direct intermediate in retrotransposition (Martin 1991; Hohjoh and F. Singer 1996; Kulpa and Moran 2005; Doucet et al. 2010; Taylor et al. 2013a, 2018). L1 insertion at the target genomic locus occurs via target-primed reverse transcription (TPRT) (Luan et al. 1993; Feng et al. 1996; Cost et al. 2002). While some key amino acid sequences have been elucidated (Mathias et al. 1991; Feng et al. 1996; Weichenrieder et al. 2004; Khazina et al. 2011; Christian et al. 2016; Ade et al. 2018; Khazina and Weichenrieder 2018), there is still much more that remains to be understood about the various L1 protein motifs and how they contribute to the L1 life cycle.

ORF1p consists of an unstructured N-terminal region (NTR), followed by three structured domains (Figure 1A), including a coiled coil (a domain consisting of an extended series of heptad repeats; the human ORF1p contains 14 of these), an RNA recognition motif (RRM) domain, and a C-terminal domain (CTD). The structure of human ORF1 has been well-characterized by x-ray crystallography (Khazina et al. 2011; Khazina and Weichenrieder 2018), culminating in a near-full-length structural model used extensively in this report (Khazina and Weichenrieder 2018). The coiled coil domain causes ORF1p to trimerize (Martin and Bushman 2001; Khazina et al. 2011), and the RRM and CTD domains are jointly responsible for single-stranded RNA-binding (Januszyk et al. 2007; Khazina and Weichenrieder 2009; Khazina et al. 2011). Recent work has shown that the extended coiled coil domain structure is metastable, in particular its N-terminal half which contains a single “stammer” insertion (residues M91, E92 and L93) in one of the heptad repeats. This stammer is thought to lead to metastability of ORF1p because the distal part of the homotrimeric coiled-coil can sample a partially unstructured state that may allow ORF1p trimers to interact with one another and form higher order structures (Khazina and Weichenrieder 2018).

**Fig. 1.**
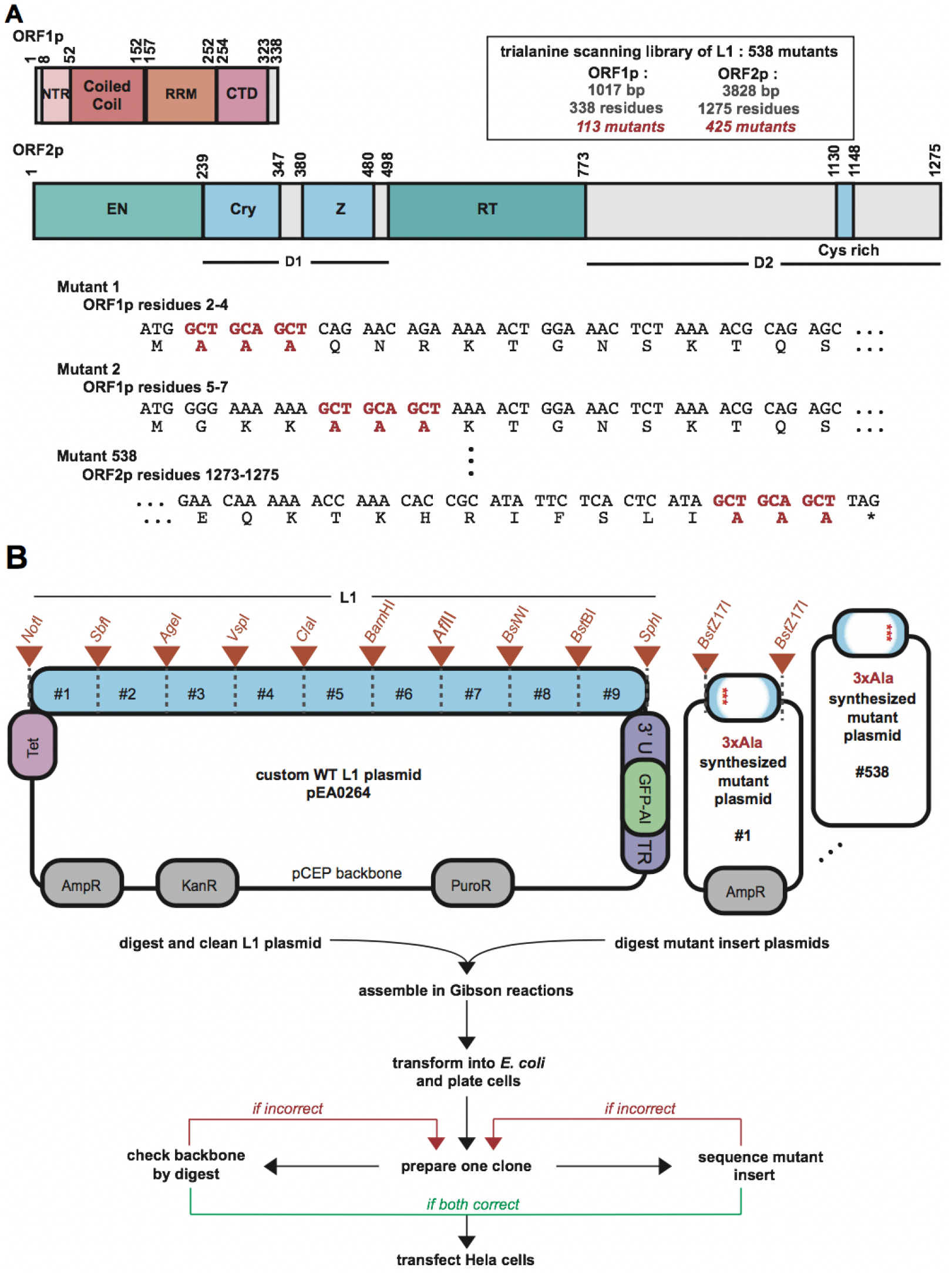
L1 architecture and the design of the trialanine scan. (A) The human L1 proteins are depicted in detail. The residue positions of characterized domains are shown for ORF1p and ORF2p. The library consists of 538 mutants. The design of the trialanine mutants for the first two and the last mutant of the library are shown at the DNA and protein sequence levels. Start and stop codons were not mutated. The trialanine mutants are consecutive and non-overlapping. (B) The parental L1 plasmid, pEA0264 is diagrammed in the upper left, featuring the engineered restriction sites. Orange triangles annotate the boundaries (designed unique restriction sites) of nine chunks. In the upper right, each of the 538 synthesized mutant plasmids were identical, excepting the 3xAla 600bp fragment provided between the BstZ17I restriction sites. The pipeline for building the library is outlined below the plasmid schematics. An efficient two-piece Gibson assembly approach followed by a two-part quality control procedure was used to build each mutant L1 construct in the library.

ORF2p also has regions of well-characterized structure and function. The most thoroughly understood regions functionally are the enzymatic endonuclease (EN) and reverse transcriptase (RT) domains (Mathias et al. 1991; Feng et al. 1996). Other less functionally defined motifs include the recently described Cryptic (Cry) sequence (Christian et al. 2016), the Z domain region (Clements and Singer 1998), and the carboxy-terminal segment (CTS), which harbors a cysteine rich motif (Fanning and Singer 1987) that is important for retrotransposition. There is a crystal structure of the EN domain (Weichenrieder et al. 2004) but the remainder of ORF2p remains structurally uncharacterized. In this work, we refer to two large, poorly characterized regions of ORF2p as Desert 1 (D1, the region between the EN and Z domains, which contains the Cry sequence) and Desert 2 (D2, the region that lies after RT and contains the CTS and cysteine rich motif) (Figure 1A).

The L1 RNP also interacts with various host-factors. RNP composition is complex and dynamic in that its intracellular location and composition changes throughout the L1 life cycle (Taylor et al. 2013a, 2018; Mita et al. 2018). Extensive research has gone into identifying and characterizing retrotransposition host factors as well as factors that influence retrotransposition (Niewiadomska et al. 2007; Beauregard et al. 2008; Suzuki et al. 2009; Arjan-Odedra et al. 2012; Dai et al. 2012; Goodier et al. 2012, 2013, Taylor et al. 2013a, 2018; Peddigari et al. 2013; Pizarro and Cristofari 2016; Liu et al. 2018). Different host factors could inhibit or facilitate L1 activity, and it is likely that ORF1 and ORF2 have coevolved with these factors. This host-specific coevolution could lead to essential amino-acid sequences that are not well-conserved.

L1 employs these endogenous activities and interactions with host factors to progress through a multi-stage life cycle. L1 RNA must be transcribed, exported, and protected from degradation. ORF1p and ORF2p must be translated, folded and co-assembled with L1 RNA. This RNP must incorporate or exclude host factors. Finally, the RNP must be imported to the nucleus, and ORF2 must mediate TPRT at a target locus. Mutating L1 affects DNA, RNA, and protein primary sequences, and thus may affect any of the steps listed above. While excellent work has begun to dissect the molecular details of this life cycle, the functional significance of most ORF1p and ORF2p residues remains unknown. Therefore, we set out to build and characterize a scanning trialanine mutant library to determine how disruption of L1 sequence may impact its cellular activities. We built 538 mutants of a human L1 and characterized this ordered library by measuring retrotransposition efficiency, ORF1 RNA and protein abundance, and ORF1p cellular localization. We also compared conservation and retrotransposition efficiency throughout ORF2p, which helped identify which areas in the poorly characterized ORF2p deserts are most interesting to study further. This first comprehensive scanning mutagenic library of any transposable element provides a map that indicates which residues are critical or dispensable for the L1 life cycle.

## Material and Methods

### Design and construction of the trialanine scanning mutagenic library

A major goal was to create a pipeline in which an ordered (as opposed to pooled) library could be efficiently assembled. The original vector backbone, extensively re-engineered in our lab, was based on the pCEP4 oriP/EBNA-based vector that replicates autonomously in primate cells (ThermoFisher pCEP4 Catalog no. V044-50), which we refer to as pCEP-puro (the original HygroR cassette was replaced with a PuroR cassette). This was the basic backbone of the parental L1-containing plasmid, pEA0264, into which each trialanine mutant was cloned (Figure 1B). We added a KanR cassette to the vector backbone to facilitate subcloning of synthetic fragments delivered in an AmpR vector. pEA0264 contained a human L1-rp cassette, expressed from the TET (inducible, minimal-CMV) promoter. The construct did not include the native L1 5’UTR sequence. The full native L1-rp 3’UTR sequence was present, and also contained the GFP-AI fluorescent retrotransposition reporter construct (Ostertag et al. 2000). Because the L1-rp native 3’UTR has a weak polyA addition signal, we also included a downstream SV40 polyA addition signal from pCEP4.

As described in the text, unique restriction sites were designed such that they fell only within L1 and not in the vector backbone and were spaced roughly equally, about every 600bp. This entailed both removing and adding (“silently”, when in a coding region) restriction enzyme cut sites from throughout the plasmid backbone and the L1-rp cassette using the GeneDesign online tool (Richardson et al. 2006). The library was optimized to facilitate downstream combinatorial cloning and manipulation of the individual mutants. The logic behind the design and the construction of the pEA0264 and the full mutant library derived from it has been extensively described in detail (Adney 2018).

The 538 trialanine mutants were generated using Gibson assembly (Gibson et al. 2009), as shown in Figure 1B. Each mutant was contained within 1 of 9 “chunks” of synthetic DNA, which effectively replaced the WT chunk. An efficient, high-throughput protocol was developed to assemble the library, perform quality control, and prepare tissue culture grade DNA for subsequent experiments (Supplemental Figures 1 and 2; Adney 2018).

### 96-well retrotransposition assay

Retrotransposition was measured as outlined in Supplemental Figure 3 using HeLa-M2 cells (Hampf and Gossen 2007). The protocol for the following has been described in detail (Adney 2018), but in brief: on day one, 25,000 HeLa cells were seeded per well in 50 μL DMEM in a 96-well plate and transfected with 60 ng DNA approximately an hour later. On day two, puromycin (puro) was added to each well to select for cells containing plasmid, and on day three, the cells were split to a black-walled 96-well tissue culture plate and doxycycline (dox) was added to induce expression of the L1 cassette, and on day 6, the cells were fixed and stained for analysis. The plates were imaged at the NYU High Throughput Biology Laboratory for data analysis, discussed below. Supplemental Figure 3 also shows controls done to prove the robustness and reproducibility of this technique.

### Quantification of retrotransposition

96-well black imaging plates (Corning product 3603) were imaged on an Arrayscan VTI using the following parameters: 5x magnification, 2-2 binning, 4 fields per well. Image analysis was performed using the Target Activation Bioapplication (Thermo Scientific Cellomics Scan version 6.6.0, build 8153). DAPI positive nuclei were identified using a dynamic isodata thresholding algorithm after minimal background subtraction. DAPI positive objects were used to identify cell nuclei and to delineate nuclear borders. A ‘circle’ (x = 2 μm) greater than the nuclear border was drawn for each cell and the GFP expression within this area was quantified. Cells expressing cytoplasmic GFP represented retro-transposition positive cells (since the limits of fluorescence were set so that no cells were considered positive for preparations of control cells lacking GFP). The reported parameters are explained as follows: Total = total number of DAPI nuclei counted; GFP+ = above GFP threshold; (*GFP* + /*Total* * 100)_*mutant*_/(*GFP* + /*Total* * 100)_*WT*_ = retrotransposition efficiency

### Statistical analysis of retrotransposition frequency

Once all retrotransposition efficiency data were acquired, we set thresholds for which trialanine mutants had a “strong effect” (depleting activity) and which had “wild-type activity” (WT). First, to set the lower threshold, we looked at mutants containing ORF2p residues known to be critical for retro-transposition and thought to be catalytic (N13, E43, D145, D205, H230 and D702), which all showed a strong effect with retrotransposition frequencies <20% of WT, providing a good calibration of the lowest activity category. By setting a conservative threshold at 25% we allowed for some biological variation in any given mutant’s inter-experimental variation in retrotransposition level.

Second, we did a statistical analysis to set the range of WT, which meant taking all the data into consideration and establishing what we did not consider to significantly deviate from WT activity (100%). We first made sure that we did not see any major batch effects between experiments; none were noted. When the data was divided into four groups based on their activity with an equal number of mutants in each group, as expected, the error decreased as the median increased. We estimated the error distribution for different number of replicates in the 4 regions by randomly resampling the data points with replacement. Using the error distribution for the group with the highest activity that contained the WT data points, we estimated a confidence interval for what represented WT activity. For mutants with 4 replicate measurements, the 99% confidence interval was estimated at 78% – 126% of the reference WT plasmid’s activity, and we use 80% as a conservative lower limit for WT activity. No mutant’s activity averaged over 125%, indicating that we did not isolate any strong “gain of function” mutant in this library.

### Immunoblot assays and statistical analysis of ORF1p of mutant protein abundance

Each ORF1p mutant was tested for protein production in two separate biological replicates. The HeLa-M2 cells were treated and harvested in a 6-well plate format and protein was extracted and measured by quantification on a Western blot, as previously described (Adney, 2018). First, all measurements were normalized by adjusting for the expression of ORF1 protein endogenously in HeLa-M2 cells (13% of the ORF1p signal in cells expressing pEA0264 WT ORF1p in these experiments is endogenous). Based on a statistical analysis of each ORF1p mutant’s protein abundance, computed in the same manner as described above for the retro-transposition activity thresholds, the 99% confidence interval estimated 50% of WT protein abundance as the lower limit. Hence, the protein levels for each mutant are referred to as either “high”, which refers to wild-type ORF1p abundance, or “low”, which refers to a protein abundance that was less than 50% that of WT (significantly depleted).

### Measurement of total RNA abundance

The RNA level of an ORF1p mutant was calculated by comparing the total RNA to the total plasmid DNA for a given ORF1p trialanine mutant and then normalizing that to the WT value. For these measurements, we took a pooled approach in which we transfected anywhere from one to fourteen mutants into one well of cells. Cell lysate was prepared from transfected HeLa cells and the total plasmid for DNA sequencing or total RNA and for RNA sequencing were isolated and the respective libraries were prepared and sequenced as described (Adney, 2018), using 36bp paired end reads on an Illumina NextSeq 500. For analysis of pooled samples, we designed a custom series of L1 reference sequences corresponding to each L1 trialanine mutant. The references were designed for each mutant: (1) with the mutant sequence (9bp) located at the center of a 75bp sequence (with 35 bp of WT L1 on either side) and (2) the exact same sequence that was fully WT. The 36 bp reads only required 1 bp of overlap with the mutant sequence to map well. We then compared read counts, as previously described (Adney, 2018).

### Quantification of ORF1p cellular localization

Transfected HeLa-M2 cells were prepared for in a 96-well plate, fixed, and stained (with the anti-ORF1p antibody, the nucleolus using an anti-fibrillarin antibody, and Hoechst 33342) for imaging analysis as described (Adney, 2018). Images were obtained using an Andor Yokogawa CSU-X confocal spinning disk on a Nikon TI Eclipse microscope and fluorescence was recorded with an sCMOS Prime 95B camera (Photometrics) with a 100x objective (pixel size: 0.11 um). 5 random fields of view were imaged per construct per experiment. One DAPI image and a 6-step 6-um Z-stack in the ORF1p channel were acquired for each field of view. Images were acquired using Nikon Elements software and analyzed using ImageJ/Fiji. Each channel was z-projected using “Sum Slices”. The data were blinded and manually scored for nucleolar localization by a naïve investigator who recorded the number of nuclei in the image, the number of nuclei that had the nucleolar phenotype, and the approximate nucleolar-to-cytoplasmic ORF1p intensity ratio of the positive cells. Nucleolar phenotype was qualitatively evaluated by normalizing a given cell’s nucleolar ORF1p intensity to its cytoplasmic ORF1p intensity and comparing it to the same ratio in cells transfected with the wild-type construct. Nucleoli were identified by DAPI and were confirmed by fibrillarin immunofluorescence in a subset of experiments. The frequency of the nucleolar phenotype was evaluated over at least 20 cells per construct. A given mutant was considered positive for the nucleolar phenotype if its phenotype rate was greater than 1 SD above the mean phenotype rate across all constructs tested.

### Generation of alignments to evaluate conservation in ORF2p

ORF2p protein sequences were translated and aligned from a compilation of L1 nucleotide sequences (Boissinot and Sookdeo 2016), the IDs of which are listed in Supplemental Table 7. Fifty-five of the sequences, including L1-rp ORF2p, were run through multiple sequence alignment analysis, followed by measurements of percent identity using Geneious (v 11.1.2; Build 2018-03-01 15:52; Java Version 1.8.0 162-b12 64 bit: Restricted R11 license). An alignment of a representative subset of these sequences is presented in Supplemental Figure 5. The program produced the percent identity score at each residue. Since we are working with three residue windows, we used the percent identity value corresponding to the residue with the highest identity score for each trialanine mutant. We binned the identify score quantities into four bins, spanning 0-29%, 30-69%, 70-99%, and 100%. We then compared these categories to the three bins of retrotransposition efficiency explained in the text (no retrotransposition, reduced retrotransposition, and WT levels of retrotransposition). Then, the status of each mutant by each of these two measures was analyzed.

## Results and Discussion

### Retrotransposition efficiency is extremely sensitive to ORF mutations

To determine amino acid sequences in ORF1p and ORF2p that are critical for L1 function, we undertook a scanning mutagenesis study, producing a library of 538 trialanine mutants scanning human L1. These L1 proteins, consist of 338 and 1275 residues, respectively (Figure 1A). To obtain a complete mutagenic scan of the ORFs, we designed an ordered library of 113 mutants for ORF1p and 425 mutants for ORF2p, totaling 538 mutants, each of which had three consecutive residues mutated to alanine (each referred to as a trialanine mutant). The mutants tiled through the proteins, did not overlap, and did not include start or stop codons (Figure 1A, Supplemental Table 1). The identities of the final constructs that made up the library are detailed in the first column of Supplemental Table 2.

We used human L1 sequence (L1-rp, accession number AF148856), derived from a retinitis pigmentosa patient cell line, that is known to be retrotransposition competent (Kimberland et al. 1999). We used the non-endogenous, doxycycline (dox)-inducible Tet-minimal CMV reporter to drive L1 expression in place of the 5’UTR-promoter sequence (O’Donnell et al. 2013; Taylor et al. 2013b). We tested the ability of each mutant to retrotranspose using a retrotransposition assay (Supplemental Figure 3A), which is the most stringent test for function; any aspect of the L1 life cycle that is impacted by our mutations should be evident. Retrotransposition efficiency values are listed in Supplemental Table 2; Figure 2 summarizes the retrotransposition efficiency of each mutant relative to wild-type and maps this value along the length of the ORFs, highlighting key motifs and previously studied essential residues.

**Fig. 2.**
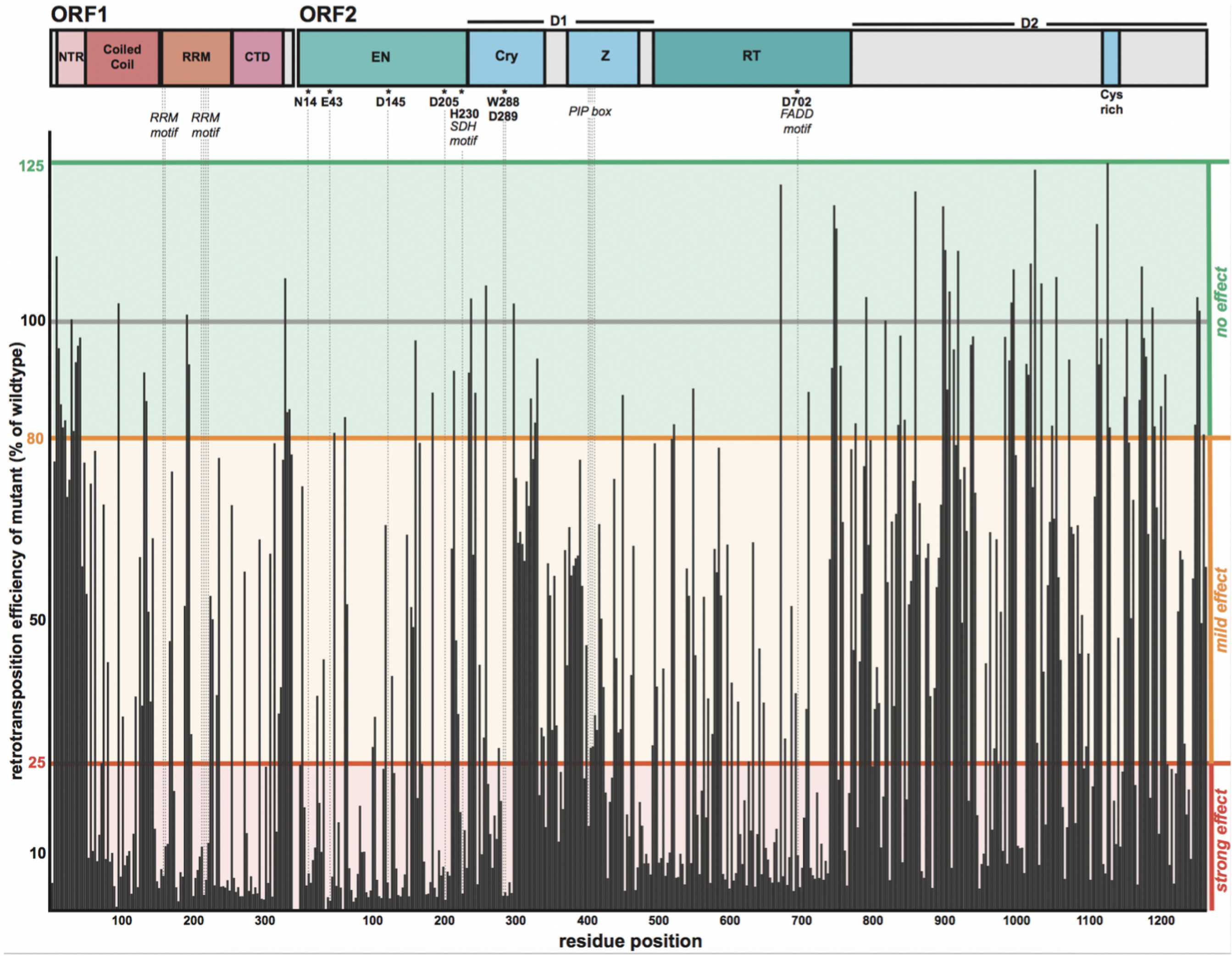
The retrotransposition efficiency of each trialanine mutant. Along the top are the schematics of ORF1p and ORF2p, highlighting domain boundaries as well as well characterized motifs and essential residues. The residue position is indicated along the x-axis. The y-axis denotes the percentage of WT activity of each mutant. Each mutant’s retrotransposition was normalized to WT measurements made in the same experiment on the same plate. WT retrotransposition frequency was set to 100% (gray bar). Statistically, values ranging between 80 and 125%were within the WT range of activity, in which the trialanine mutation had *no effect* (green background). A mutant was classified as *mild effect* for values ranging between from >25 and <80% (orange background) and *strong effect* for values of 25% and below (red background).

**Table 1.** Impact on retrotransposition efficiency organized by protein domain. The percentages of 3xAla mutants showing a strong, mild, or no effect on retrotransposition efficiency are represented for both ORF1p and ORF2p. The values for the full-length protein and then for each domain are shown.

| % of 3xAla mutations in each retrotransposition efficiency category: |  |  |  |
| --- | --- | --- | --- |
| ORF1p |  |  |  |
|  | strong | mild | none |
| Full-length Protein | 53 | 31 | 16 |
| NTR | 6 | 35 | 59 |
| Coiled coil | 56 | 35 | 9 |
| RRM | 70 | 24 | 6 |
| CTD | 55 | 32 | 13 |
| ORF2p |  |  |  |
|  | strong | mild | none |
| Full-length Protein | 48 | 35 | 17 |
| EN | 74 | 19 | 8 |
| D1 | 38 | 47 | 15 |
| Z | 36 | 61 | 3 |
| RT | 62 | 28 | 10 |
| D2 | 32 | 41 | 27 |

Retrotransposition efficiency was extremely sensitive to ORF1p and ORF2p mutations, consistent with expectations for a “streamlined” and highly conserved element. About 50% of the trialanine mutants had a strong effect, 34% had a mild effect, and only 16% retained wild-type activity. A significant fraction of the total mutants (25% of ORF1p and 12% of ORF2p mutants) had activity <=5% of WT. None of the mutants caused a significant increase in activity (>125% of wild-type). ORF1p and ORF2p had similar frequencies of deleterious mutations, with obvious clusters of strong effect in the more conserved domains of the proteins (Table 1).

Overlaying retrotransposition levels of the mutants on solved WT crystal structures gives a visual representation of each mutant’s impact, for example the EN domain of ORF2p (Supplemental Figure 4). The mapping of mutant phenotypes onto the full-length ORF1p structure model will be presented visually in the next section, together with protein abundance data.

Mobile elements that remain active in the human genome inspire comparison to host-parasite arms races (Daugherty and Malik 2012). While L1 is not simply a parasite and does play important roles, L1 elements also pose a strong risk to the host due to their strong mutagenic capacity and so the element can be considered to be analogous to a parasite with respect to the evolution of its DNA sequence. The host is likely under strong selection to reduce retrotransposition while L1 must evolve a robust life cycle to avoid extinction. This type of antagonistic selection tends to minimize genome sizes in obligate-parasitic organisms (Wolf and Koonin 2013). In addition, the biochemistry of the L1 life cycle may drive genome minimization. The huge number of truncated L1 remnants in the human genome suggests that the reverse transcriptase step is frequently not processive enough to drive successful retrotransposition in the host-environment. This may be an intrinsic limitation of the RT enzyme, but it is also likely that the host has evolved mechanisms that actively promote 5’ truncation. Thus, shortening the L1 genome would increase its probability of propagation. The net result would be an increase in information density in the protein coding regions in the element. The high density of critical regions for retrotransposition that we found provides strong evidence for this streamlining hypothesis.

### Most ORF1 mutants are expressed robustly

We quantified the relative protein levels of the ORF1p mutants individually by immunoblotting (Figure 3A and Supplemental Table 3) using a monoclonal antibody targeting endogenous human ORF1 (Rodić et al. 2014). Due to substantial variations (2-fold) in ORF1p levels in replicate immunoblot experiments we treated the average protein abundance for each mutant as binary, with a conservative cut off: high (>50% that of WT) or low (<50%). Only 24% of the mutants (27/113) resulted in ORF1p reduction to <50% that of wild-type. All of these mutants with low ORF1p also showed loss of retrotransposition activity. Retrotransposition and protein abundance data are summarized in Supplemental Table 4. Trialanine mutants that disrupt the epitope that our antibody recognizes could not be assessed by Western blot. However, since all these mutants showed WT or close-to WT levels of retrotransposition, we can confidently surmise that they were well expressed. Figure 4 summarizes the effects of protein levels and retrotransposition activity mapped onto along the ORF1p crystal structure. Of the mutants that show low protein levels, all map to the RRM and CTD domains (16 and 11 mutants, respectively; Figure 3B). We speculate that these mutants interfere with the folding of these highly structured domains.

**Fig. 3.**
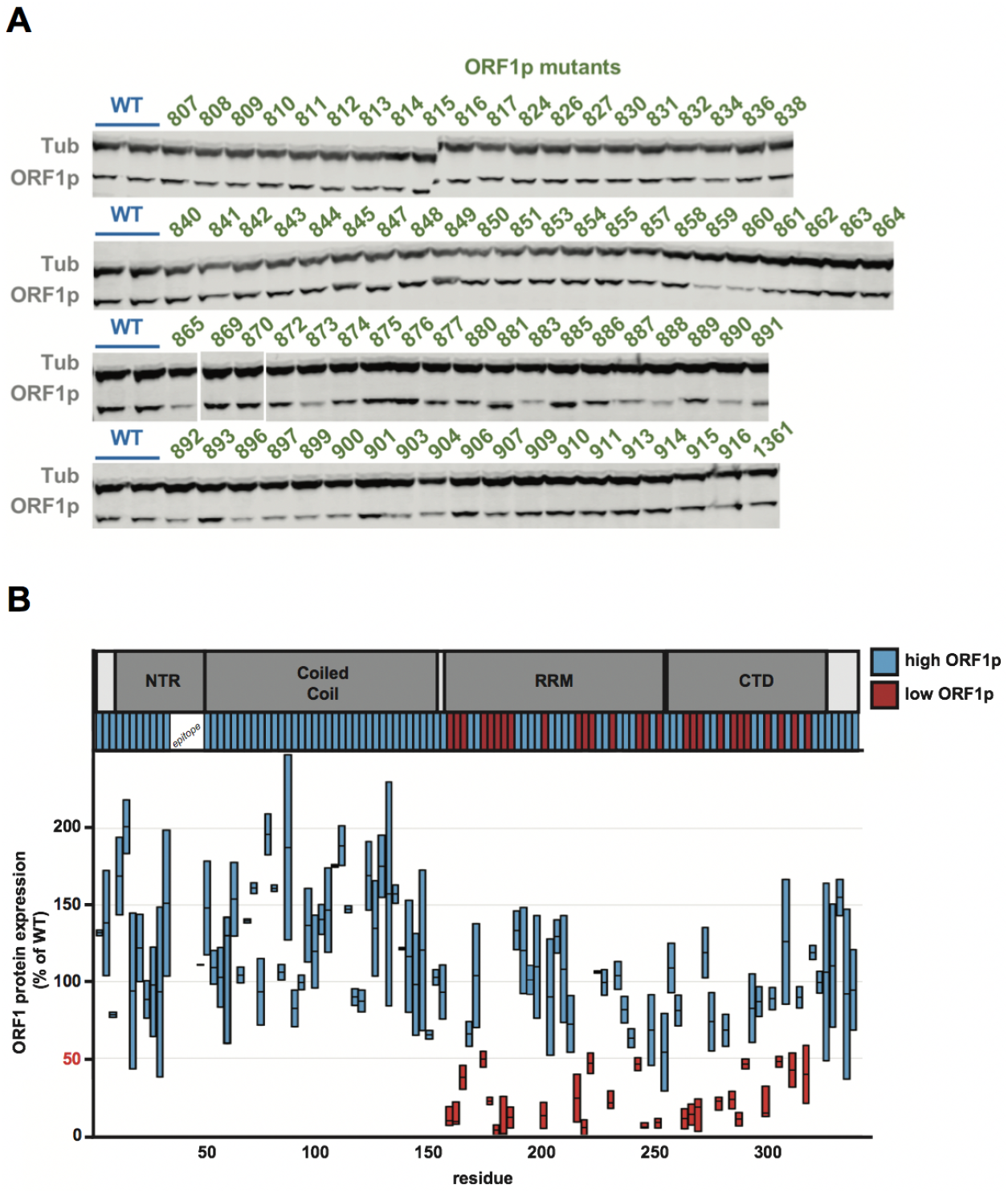
Protein abundance of mutants of ORF1p. (A) Representative immunoblots for WT pEA0264 and the ORF1p mutants. Samples were prepared from 6-well plates of HeLa cells, the clarified lysates of which were probed with anti-ORF1 and anti-tubulin antibodies. HeLa cells lacking a plasmid reproducibly expressed ORF1p at a level of 13% of pEA0264. (B) The ORF1p schematic is shown atthetop. Resultsfrom immunoblot analysesfor each ORF1p mutantare represented on the plot. Two measurements are shown for each mutant, quantified from independent experiments. These values were background subtracted to remove signal corresponding to endogenous ORF1p expression. Protein levels are plotted on the Y axis and residue position is indicated on the X-axis. We observed some variability and thus plotted the range for each mutant as a bar with a horizontal bar marking the mean. We refer to protein abundance in binary terms, as either high (+) or low (-), using 50% (marked in red) as the threshold. The mutants are color coded in the bar below the ORF1p schematic to highlight which regions had high (blue) or low (red) protein levels.

**Fig. 4.**
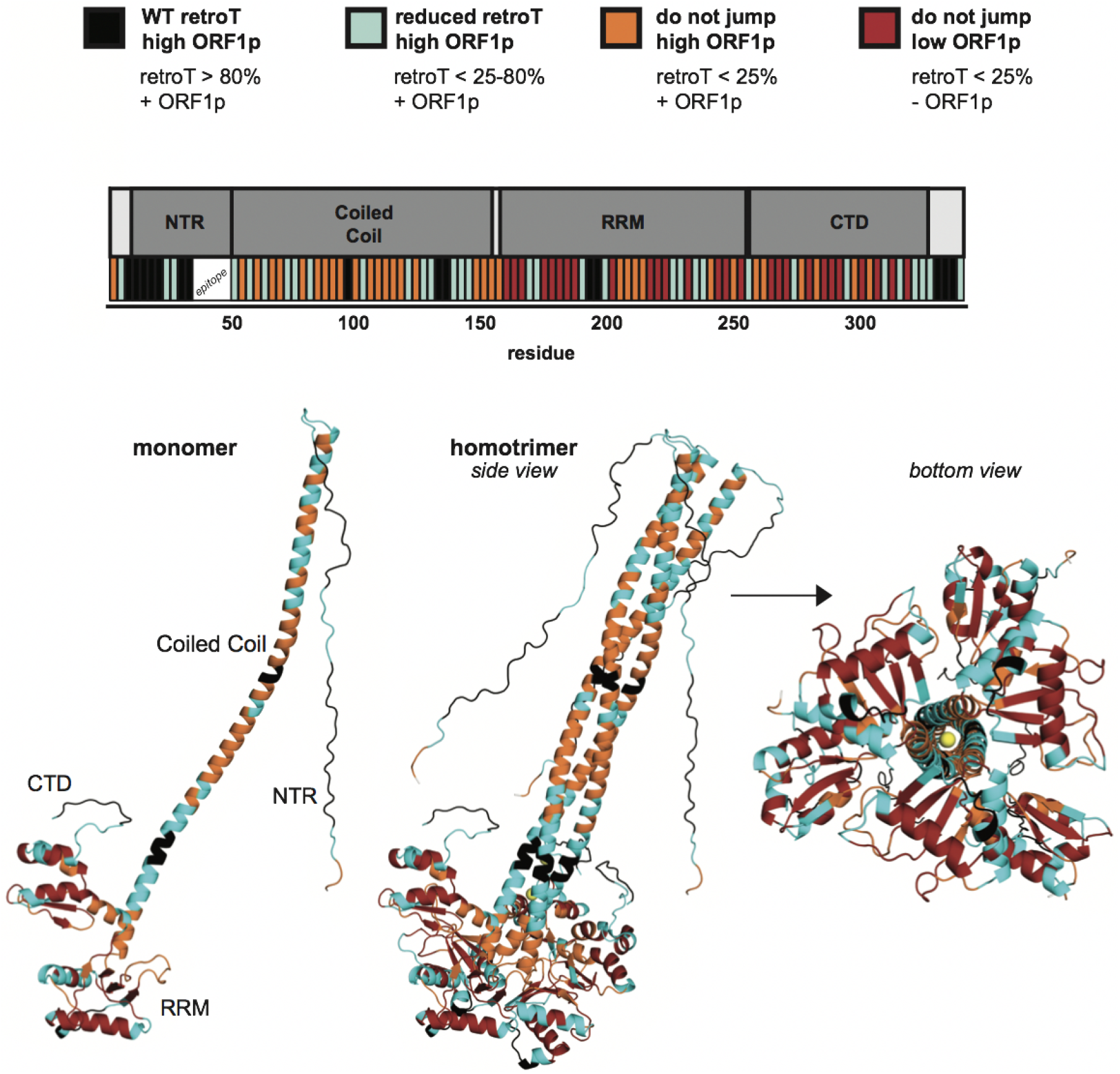
Retrotransposition and protein abundance of mutants mapped onto a three-dimensional model of trimeric ORF1p. The model is based on available crystal structures (Khazina and Weichenrieder 2018). The mutants are divided into four categories and color coded, shown at the top. This provides a visual representation of retrotransposition efficiency and protein abundance, both along the linear schematic of ORF1 with the corresponding color-coded bars as well as projected onto the WT ORF1p monomer and trimer structures. The color code is as follows: high ORF1p and WT retrotransposition (black), high ORF1p and reduced retrotransposition (cyan), high ORF1p and no retrotransposition (orange), low ORF1p and no retrotransposition (red), chloride ions noted in the structure of (Khazina et al. 2011) (yellow), and the initial methionine (not mutated, white).

Next we wished to evaluate the effect of each mutant on L1 RNA stability. To do so, we designed pooled RNA-seq and DNA-seq experiment to evaluate the impact of ORF1p mutations on RNA abundance. The experimental design is shown in Figure 5A-B: hypothetical Mut X has abundant RNA is abundant while Mut Y has low abundance RNA. DNA-seq reads were used to normalize the transfection efficiency of each plasmid. Sequencing reads containing the unique 9 bp of each 3xAla insertion were used to determine RNA and DNA levels. The WT plasmid and RNA were used as internal controls for WT L1 behavior. In this way, we determined the relative RNA abundance every mutant.

**Fig. 5.**
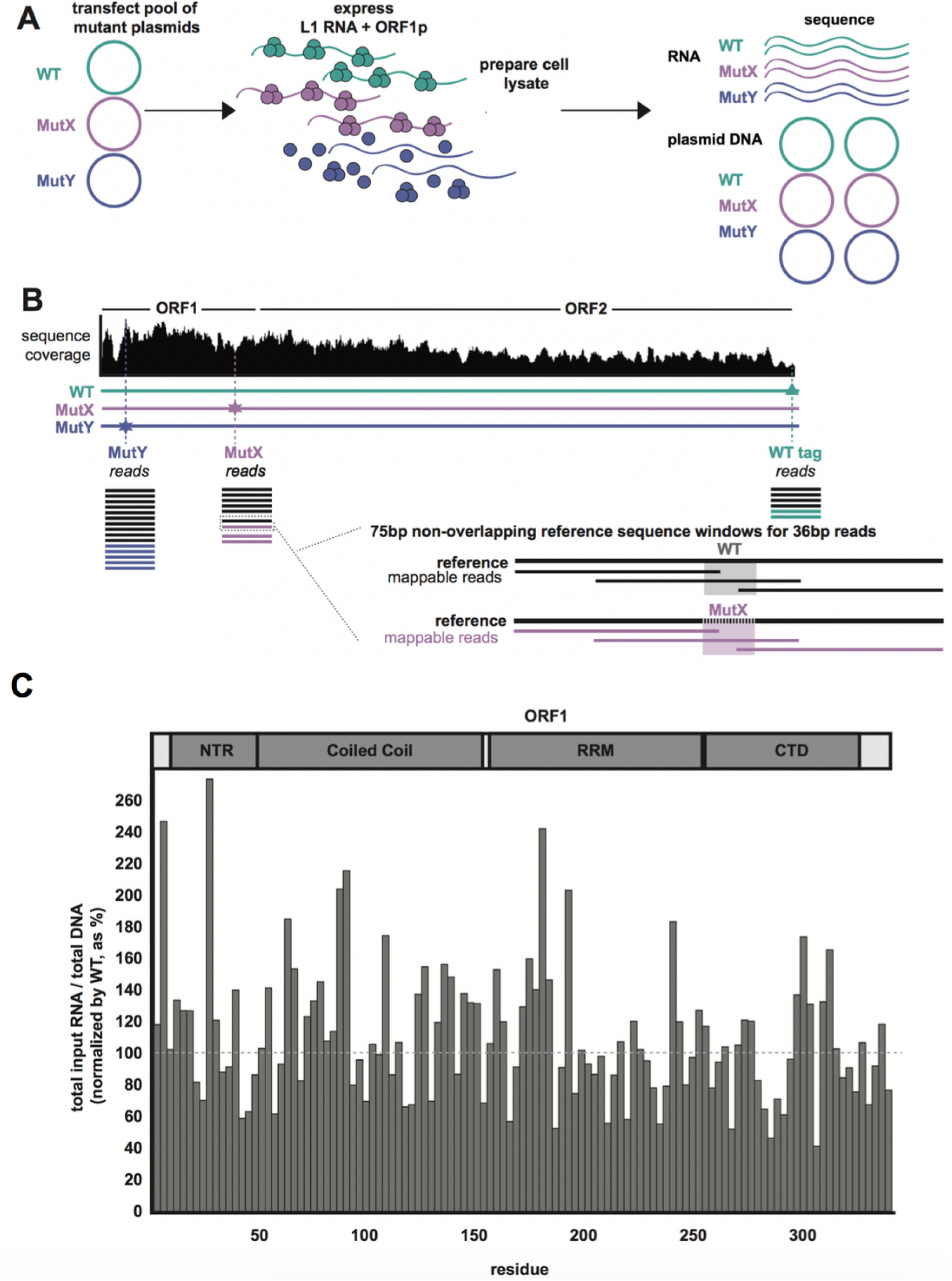
The protocol for a pooled approach to sequence plasmid DNA and total RNA of L1 mutants. (A) The workflow for transfecting a pool of two mutants and the WT plasmids (thus co-expressing three constructs), preparing cell lysate, and sequencing the isolated pools of L1 plasmid DNA and total RNA is shown. In this depiction, at the end of the experiment, all three plasmids have an equal abundance of plasmid DNA copies. Mut X and WT show equal L1 RNA abundance, while that of Mut Y is reduced 5-fold. (B) Sequence coverage across L1 coding region is uneven. This diagram depicts how reads were mapped and was normalized to both the sequencing depth at a given window as well as to the internal WT plasmid control. (C) The RNA abundance (normalized to plasmid DNA abundance) of each ORF1p mutant is shown as the percent of WT. The fraction of total mutant L1 RNA in the lysate is shown, normalized to the WT level. The gray dashed line indicates WT levels at 100%.

We pooled several mutants at once with the WT construct for a total of 8 pools (named Pool 1-Pool 8; Supplemental Table 5) and expressed each of them in human cells. The data are reported as RNA abundance of each mutant in Figure 5C. Notably all mutants had near-WT levels of RNA abundance, and no mutant had <60% that of WT, indicating that RNA abundance (reflecting transcription efficiency and stability) is unlikely to explain ablation of retrotransposition in many of the ORF1p mutants. However, a formal demonstration of this will require replicates, an expensive experiment for what is likely to be a negative result.

### Two coiled-coil mutants have significant relocalization to the nucleolus

We evaluated whether any ORF1p mutants that block retro-transposition might do so by interfering with proper subcellular localization. Nucleocytoplasmic trafficking is key to the L1 life cycle and our previous studies revealed relocalization of ORF1p from the cytoplasm to the nucleus during the M/G1 phase of the cell cycle (Mita et al. 2018). We therefore used immunofluorescence (IF) to probe the localization of the 40 ORF1p trialanine mutants that produce normal levels of protein but have decreased retrotransposition activity (Supplemental Table 6). The vast majority of the ORF1p mutants localized primarily to the cytoplasm, just like WT ORF1p. However, we observed that two ORF1p mutants, CLK86-88AAA and LRS107-109AAA, displayed strong nucleolar localization in a subset of cells (Figure 6 and Supplemental Table 6). This striking relocalization phenotype was seen in 8% and 24% of total cells for the CLK86-88AAA and LRS107-109AAA mutants, respectively, as compared to <1% of cells expressing WT ORF1p. We did not observe a correlation between nucleolar localization and total ORF1p fluorescence in a given cell.

**Fig. 6.**
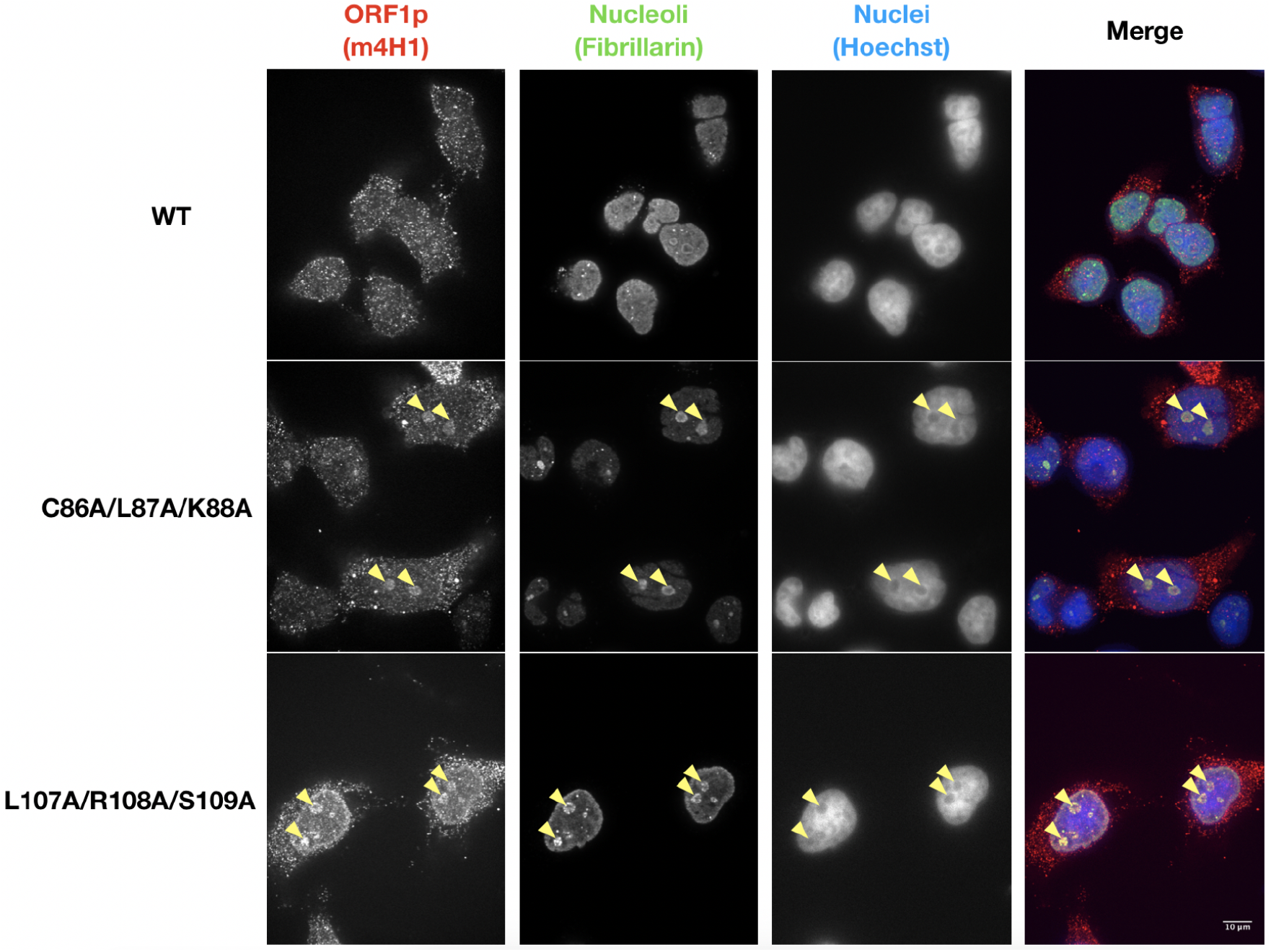
Immunofluorescence analysis reveals intriguing nucleolar localization of a small subset of ORF1p mutants. Representative images of immunostained HeLa-M2 cells expressing wild-type (WT) L1 (top) or L1 ORF1p mutants (CLK86-88AAA *[middle]* and LRS107-109AAA [bottom]). Yellow arrowheads indicate nucleoli with diffuse ORF1p localization in the mutant construct transfections. Cells were stained with mouse anti-ORF1p *(left*), rabbit anti-fibrillarin *(middle left*), and Hoechst 33342 *(middle right)*. Antibody target names are reported above the corresponding pictures and colored according to the colors used in the merged pictures *(right)*. Scale bar = 10 um.

Both mutants of interest reside in the coiled coil domain of ORF1p. A C86S substitution was previously shown to strongly reduce retrotransposition, which was surprising given the poor conservation of C86 across primate L1 sequences and its position on the surface of the coiled coil (Khazina and Weichenrieder 2018). Our data with CLK86-88AAA recapitulates the sharp decrease in retrotransposition and suggests a defect in intracellular localization as a potential mechanism. Additionally, a three-residue insertion in the stammer portion of the heptad repeat structure of the ORF1p coiled coil (residues 91 – 93) was proposed to contribute to the structural malleability of the coiled coil N-terminal to the stammer (Khazina and Weichenrieder 2018). They proposed a model in which the stammer introduces flexibility into the coiled coil that then allows for ORF1p trimers to adopt an open conformation and form inter-trimer interactions between ORF1p N-termini. These inter-trimer interactions were suggested to drive higher-order ORF1p structures, such as linear arrays and a larger meshwork of trimers. The stammer lies between our two trialanine mutants of interest. It is conceivable that the CLK86-88AAA and LRS107-109AAA mutants change the flexibility of the ORF1p coiled coil in similar ways, thus interfering with the L1 life cycle and increasing the propensity of the protein to localize to the nucleolus. While the reasons for nucleolar localization will require further investigation, we speculate that the localization of a subset of ORF1p mutants to the nucleolus could be the result of altered binding affinities for nucleic acid or protein partners.

Previous work on LINE-1 proteins and the nucleolus demonstrated localization of WT ORF1p to the nucleolus in close to 50% of 143B TK cells (Goodier et al. 2004). However, this localization was tag-dependent and was seen either in ORF1p-only expression constructs or in bicistronic constructs with two IRESs, which complicates interpretation. Further exploration of ORF1p-only constructs identified a E165G ORF1p mutant that has enhanced nucleolar localization and also indicated that nucleolar localization is likely RRM-dependent since actinomycin D treatment abolished nucleolar localization of WT and E165G ORF1p without changing cytoplasmic foci formation (Goodier et al, 2007). Taken together, we expect that ORF1p localization to the nucleolus might be a physiological step in the L1 life cycle but more likely that accumulation of ORF1p in the nucleolus may instead be a cause or consequence of L1 transposition defects. Notably, other work showed that the L1 RNA itself interacts with nucleolin, a nucleolar protein, and promotes transcriptional program changes that are necessary for embryonic development in mice (Percharde et al. 2018). However, while L1 RNA was predominantly nuclear in these mouse embry onic stem cells (mESCs), ORF1p was mostly cytoplasmic. It is possible that in our cell system, endogenous nucleolin captures more ORF1p-bound L1 RNA and that ORF1p muta tions alter the ability of nucleolin to bind to ORF1p-decorated L1 RNA. Interestingly, nucleolin was previously identified as a factor that specifically promotes ORF2p translation, and nucleolin knockdown was found to decrease L1 retrotransposition rates (Peddigari et al. 2013). Thus, localization of ORF1p mutants to the nucleolus may be indicative of an imbalance of L1 RNA interactions with ORF1p and nucleolin, which could in turn lead to a decrease in L1 retrotransposition rates.

### A cluster of transposition-defective mutations in a nonconserved domain of ORF2

Similar to ORF1p, where some of the least conserved portions of the protein are functionally essential (Khazina and Weichenrieder 2018), there could also be such regions in ORF2p, which might not be detectable purely through sequence analysis, but only using functional analysis. We therefore not only mapped retrotransposition efficiency onto the crystallized endonuclease domain of L1 ORF2p (Supplemental Figure 4), but also correlated retrotransposition efficiency with sequence conservation all along the ORF2p sequence. To this aim we aligned the human ORF2 protein sequence to 14 diverse mammalian sequences as well as others from more distant vertebrates (Supplemental Table 7 and Supplemental Figure 5). As expected we found highly conserved sequences to be important for retrotransposition. Importantly, however, we also identified clusters of functionally crucial residues in the less conserved regions.

Until now, conservation of functional residues summarized as short amino acid sequence motifs, has been integral to identifying regions of ORF2p indispensable for L1 activity. However, there are “desert” regions (D1 and D2) in ORF2p that have no structural motifs and no clear conservation. Our unbiased scanning approach helps us reach beyond the most studied regions of ORF2p and creates a framework for prioritizing functional regions for further study. Figure 7 summarizes both the conservation and retrotransposition frequencies of each ORF2p mutant. A few previously noted amino acid sequence motifs were confirmed to be essential by this analysis, such as the “Cry motif’ in the D1 region and the Cys rich motifs in the D2 region. However, there were also regions that lacked amino acid sequence conservation but showed a profound retrotransposition defect. We denoted these positions with stars in Figure 7. This analysis revealed a “Star Cluster”, contained in the window of residues F952 – C1020, a previously uninvestigated region with a high density of amino acid sequences of this type. This region is now of special interest for further characterization.

**Fig. 7.**
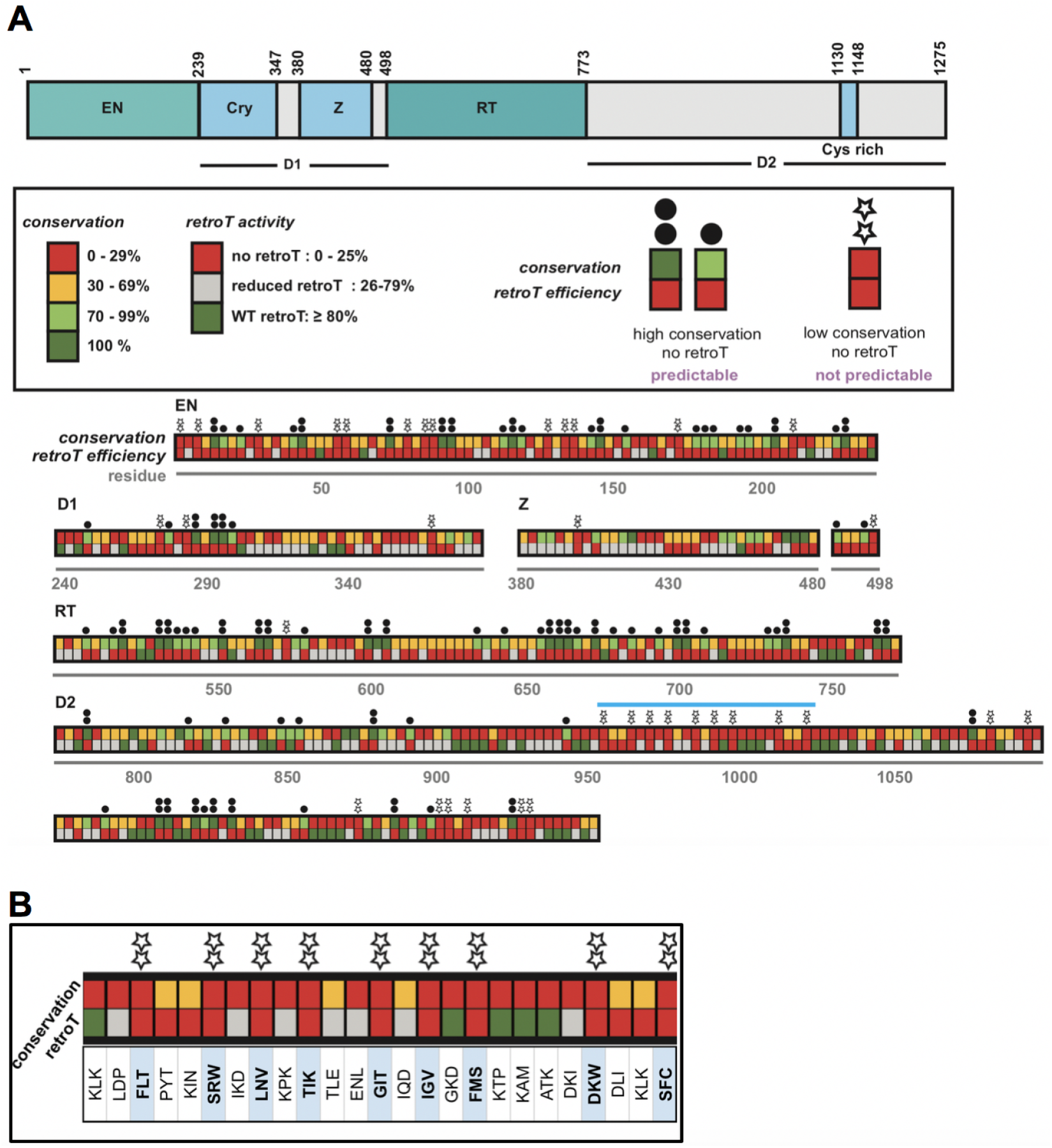
Trends in amino acid conservation and sensitivity to mutation across ORF2p. (A) The schematic for the ORF2p domains is along the top. This is a graphical representation and interpretation of the conservation data displayed in Supplemental Table 8 (mammals and all vertebrates; the ‘mammals alone’ column is excluded). For each trialanine mutant, we took the value corresponding to the residue with the highest conservation to represent the mutant. As shown in the box, conservation and retrotransposition are color-coded. The boxes are stacked to compare conservation and activity. One dot and two black dots above a mutant mean that, as expected, there was strong conservation and no L1 jumping. Stars above the mutant mean that there is low conservation and no retrotransposition, highlighting areas that may be important in ORF2p not predicted by conservation alone. The *Star Cluster* region is indicated with a light blue bar, which is shown (B) zoomed in and in detail with the three WT amino acids (in single letter format) corresponding to each mutant.

We report here the most comprehensive ordered and arrayed amino acid substitution library for any retrotransposon, DNA transposon or retrovirus. We anticipate that this resource will be of substantial interest to students of these elements and may serve as a model for future libraries of this type.

## ACKNOWLEDGEMENTS

This work was supported in part by NIH grants P50 GM107632 to J.D.B. and P01 AG051449 to John Sedivy and J.D.B. We thank Elena Khazina and Oliver Weichenrieder for the structural coordinates of their composite L1ORF1p model and for sharing information before publication. We thank Zoltán Ivics for support of M.T.O. during his visit to our laboratory. We also thank Kathleen Burns (reader), David Graham, Jeremy Nathans, and Roger Reeves for input throughout the project and for serving on the PhD thesis committee of E.M.A.

