## SupplementalMaterial for "Comprehensive scanning mutagenesis of human retrotransposon LINE-1 identifies motifs essential for function"

### Supplemental Material : Figure and Table Legends

**Supplemental Table 1** Organization of cloning trialanine mutants into nine chunks. The L1 coding sequence was divided into nine chunks. Mutants contained in each chunk are shown.

**Supplemental Figure 1** Detailed high-throughput cloning procedure used to create library. The detailed steps for building and validating the library in a high-throughput manner are depicted. The inset shows an agar plate after overnight growth of eight *E. coli* transformations of putative trialanine clones after using the "drop method".

**Supplemental Figure 2** Restriction digest used to validate final constructs in library. *Pst*I digestion was used to validate the final integrity of the backbone and the presence of the trialanine mutation for each clone. On top, a schematic of where *Pst*I cuts in the plasmid is shown. The trialanine DNA sequence was designed to contain a *Pst*I site, thus the predicted band sizes for each construct are unique. An agarose DNA gel shows diagnostic digests of each final correct clone in chunk 3, Mutants 110-160 ("WT" is the pEA0264 parent backbone and "L" is the 2-log ladder). Sanger sequencing (not shown) also confirmed the correct sequence for each mutant insert by spanning the Gibson homology arm boundaries for each clone.

**Supplemental Table 2** Raw retrotransposition efficiency data for full library. For ORF1p and ORF2p, this table shows the IDs and raw retrotransposition data for each mutant. The column *Trialanine construct ID* describes each mutant using the following format (with the information, separated by underscores): the pEA construct number, the first residue mutated, the last residue mutated, and the three WT amino acids (single letter code) mutated to alanine. The second column shows the raw data that corresponds to Figure 2: *retroT average.* The third and fourth columns show the *standard deviation* and *number of measurements*, respectively, for the measurements made for each mutant.

**Supplemental Figure 3** High-throughput microscopy-based measurement of retrotransposition. (A) The protocol for the six-day 96-well, microscopy-based assay for plasmid transfection and measurement of the retrotransposition activity in human HeLa cells. Puromycin selection of the cells containing plasmid for five days was followed by dox-induced expression of L1 for four days. (B) Representative pictures of transfected WT constructs. This photo represents one quarter of one well in a 96-well plate. This shows two channels (DAPI and GFP) and the overlay. Blue (DAPI, alive) and green (GFP, retrotransposition-positive) cells were quantified. Raw wild-type retrotransposition values were reproducible and robust, usually showing ~13% GFP+ of live cells. (C) Because DNA was purified for 538 constructs in different batches, we wanted to be sure that there was no effect of the batch in which DNA was prepared on the retrotransposition efficiency. We measured retrotransposition for WT and two independent preparations of 5 different (color coded) mutants (chosen at random). (D) Comparison of transfecting different amounts of DNA. The fact that the retrotransposition frequency is independent of DNA concentrations within the range of concentrations used here shows that small fluctuations DNA stock concentration will not affect comparisons among mutants. Cells died when transfected with too much DNA and gave reproducible results over a concentration range of 7- 68 ng (60 ng was used for experiments unless indicated otherwise).

**Supplemental Figure 4** Retrotransposition levels mapped onto EN 3D structure. The crystal structure of the WT EN domain of ORF2 (PDB 1VYB) is shown and is color-coded to display retrotransposition efficiency of each trialanine mutant. Red: strong impact on retrotransposition in red, Gray: mild impact, Cyan: no impact. (Note that this color scheme is not identical in all the figures).

**Supplemental Table 3** Raw protein expression data. Each mutant was measured for protein expression twice and the average value (after endogenous HeLa ORF1p subtraction) is shown. Gray indicates the mutants that were not measurable with this approach because the epitope was mutated. The other values are shown in blue (high ORF1p) or red (low ORF1p) and coincide with the representation in Figure 3B.

**Supplemental Table 4** Categories of ORF1p mutants based on activity and protein abundance. The percentages of ORF1p mutants that show a strong, mild, or no effect on retrotransposition efficiency combined with whether or not they impact ORF1 protein abundance are represented in 4 categories (that correspond to Figure 4) for ORF1p. The values for the full-length protein and then for each domain are shown.

**Supplemental Table 5** Composition of mutant pools for RNP studies. The composition of each of the pools (Pool 1- Pool 8), each mutant is listed using the pEA clone number as the ID. This experiment was done fully for the complete series of mutants of ORF1p, in which we measured levels of total DNA and RNA in the cell lysate.

**Supplemental Table 6** An analysis of ORF1p mutants that localize to the nucleolus. Comparison of nucleolar localization phenotype in immunocytochemistry of ORF1p mutants. 40 of the 113 ORF1p mutants in the mutant library were evaluated for nucleolar ORF1p localization by immunofluorescence staining followed by blinded counting. Two mutants were considered positive for the nucleolar phenotype, with a nucleolar phenotype rate of greater than one standard deviation above the mean nucleolar phenotype rate across all tested mutants. A minimum of 20 cells were counted for each mutant. The ORF1p domain column refers to the region of the protein in which the mutation occurs for a given mutant; NTR: N-terminal region, CC: coiled coil, RRM: RNA recognition motif, CTD: C-terminal domain. Retrotransposition rates are separated into 3 categories: high (++, > 80% of wild-type), reduced (+, between 25 and 80% of wild-type), and poor (–, < 25% of wild-type). All mutants tested produced more than 35% of wild-type ORF1p protein levels as measured by Western blot. The number of measurements indicates the number of separate experiments that include independent transfections of the given mutant.

**Supplemental Table 7** The IDs of the fifty-five LINE-1 ORF2p sequences used for phylogenetic conservation analysis. We used the human ORF2p from L1-rp (accession number AF148856). For the remainder, the DNA sequences were provided by Stephane Boissinot (Boissinot and Sookdeo 2016) and were then prepared for protein multiple sequence alignment by Oliver Weichenrieder. These were carefully curated as the most reliable mammalian and non-mammalian vertebrate ORF2p consensus sequences. (Note that these were used as the basis for analyses presented in Supplemental Table 8 and Figure 7.)

**Supplemental Figure 5** Alignment of a subset of the ORF2p sequences presented in Supplemental Table 7. This shows a multiple sequence alignment produced by Geneious for human L1-rp and 25 other L1 (13 mammalian and 12 non-mammalian vertebrate) ORF2p sequences. The identities of the organisms are along the far left (highlighted in purple). The domains of ORF2p are indicated in tracks above the alignment, which is numerated every 10 amino acids for the reference human ORF2p sequence. This used a Blosum62 Score Matrix and the individual residues are color coded as follows: 100% similar (green), 80-100% similar (greenish-yellow), 60-80% similar (yellow), <60% similar (white). Listed here are how we refer to the IDs (Supplemental Table 7) in the alignment (shown in parentheses) : HUMAN L1-rp (HUMAN) , RABBIT (RABBIT) , PIG (PIG) , COW (COW) , DOG (DOG) , PANDA (PANDA) , HORSE (HORSE) , LEMUR (LEMUR) , ELEPHANT (ELEPHANT) , HYRAX (HYRAX) , ARMADILLO (ARMADILLO) , MOUSE (MOUSE ) , RAT (RAT) , OPOSSUM (OPOSSUM) , FROG_L1-6 (FROG_6) , FROG_L1-32 (FROG_32) , FROG_L1-15 (FROG_15) , FROG_L1-29 (FROG_29) , ZEBRAFISH_L1-1 (FISH_1) , ZEBRAFISH_L1-7B (FISH_7B) , ZEBRAFISH_L1-13A (FISH_13A) , ZEBRAFISH_L1-

17B (FISH_17B) , LIZARD_L1_AC1 (LIZ_AC1) , LIZARD_L1_AC14 (LIZ_AC14) , LIZARD_L1_AC15 (LIZ_AC15) , LIZARD_L1_AC17 (LIZ_AC17).

**Supplemental Table 8** Phylogenetic conservation values of ORF2p mutants used for sensitivity analysis. The sequences shown in Supplemental Table 7 were aligned using the Geneious multiple protein sequence alignment tool and identity calculation tools. Column one indicates the ORF2p trialanine mutant. Columns two and three show the conservation results for mammals only and then for mammals and other vertebrates. Since each mutant spans three amino acid residues, the value shown is that of the residue with the highest identity score. The fourth column shows the average retrotransposition of the given mutant. The last column indicates the domain of ORF2p in which the mutant lies. For all columns, low is indicated in red and high is indicated in green. The last two columns show the retrotransposition efficiency (green : >80% of WT, gray : 25-80% of WT, and red : <25% of WT) and domain within which each mutant is located.

### Supplemental Material : Figures and Tables

**Supplemental Table 1**

| chunk # | range of mutated residues ORF1 | ORF2 | total # residues | total # 3xAla mutants | unique flanking restriction sites | |
|---|---|---|---|---|---|---|
| **1** | 2 - 175 | | 174 | 58 | *Not*I | *Sbf*I |
| **2** | 176 - 328 | | 153 | 51 | *Sbf*I | *Age*I |
| **3** | 329 - 338 | 2 - 144 | 153 | 52* | *Age*I | *Vsp*I |
| **4** | | 145 - 315 | 171 | 57 | *Vsp*I | *Cla*I |
| **5** | | 316 - 504 | 189 | 63 | *Cla*I | *Bam*HI |
| **6** | | 505 - 717 | 213 | 71 | *Bam*HI | *Afl*II |
| **7** | | 718 - 903 | 186 | 62 | *Afl*II | *Bsi*WI |
| **8** | | 904 - 1083 | 180 | 60 | *Bsi*WI | *Bst*BI |
| **9** | | 1084 - 1275 | 192 | 64 | *Bst*BI | *Sph*I |

* The final mutant for ORF1 mutates residue 338 (1xAla).
The first mutant for ORF2 mutates residues 2-3 (2xAla).

#### Supplemental Figure 1

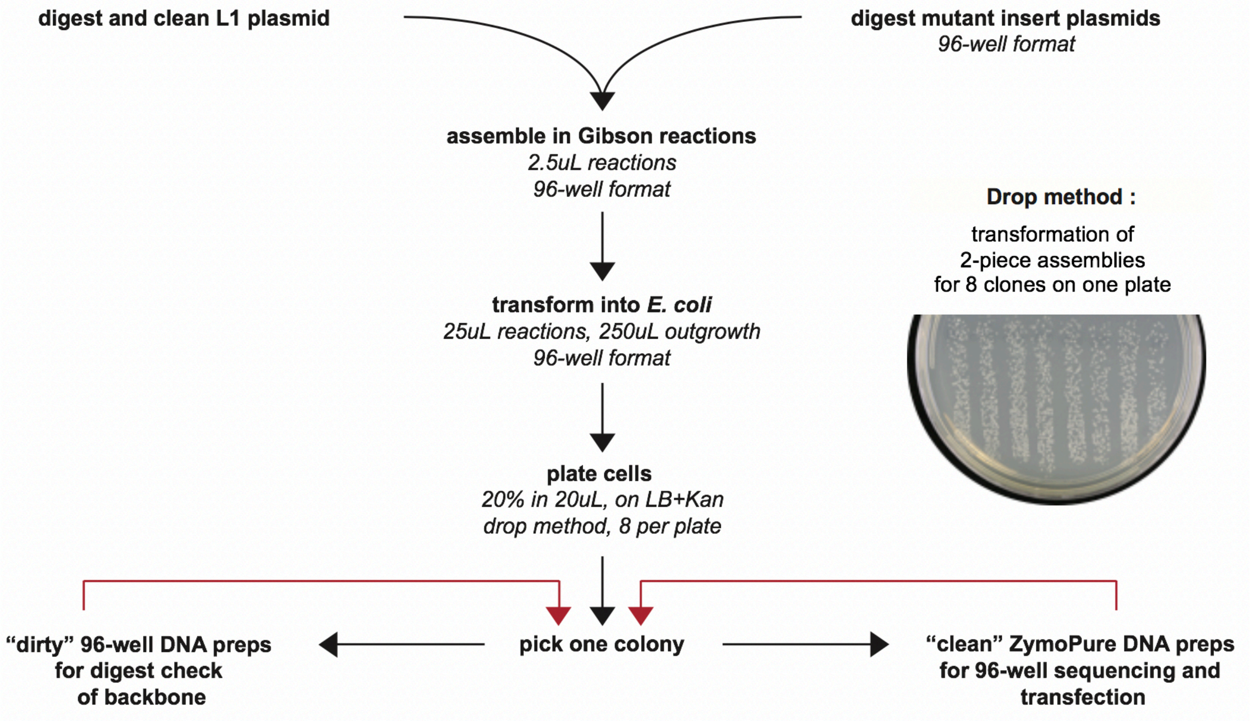

#### Supplemental Figure 2

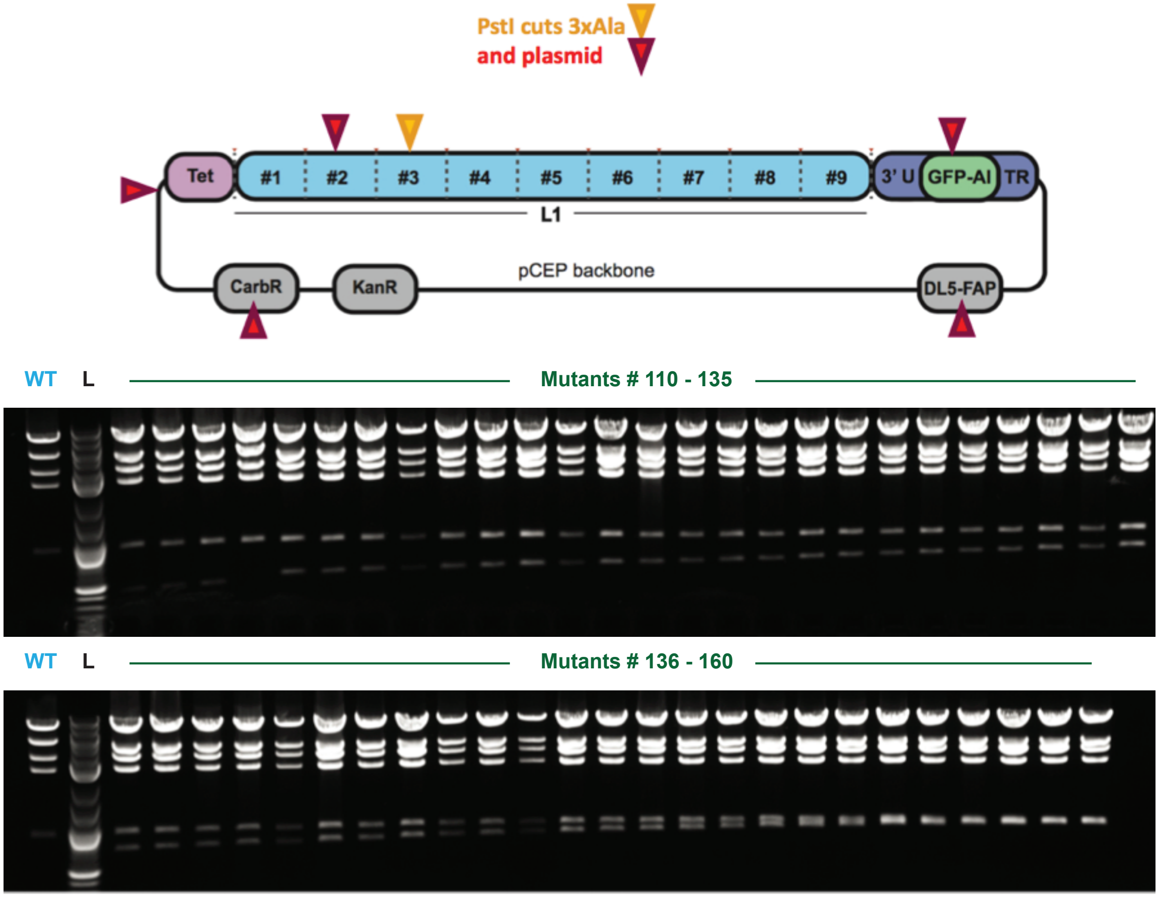

#### Supplemental Table 2

| ORF1 | | | | | | | |
|---|---|---|---|---|---|---|---|
| Mutant ID | retroT average | standard deviation | number of measure-ments | Mutant ID | retroT average | standard deviation | number of measure-ments |
| pEA807_2_4_GKK | 4.8 | 3.8 | 4 | pEA863_170_172_NGT | 74.5 | 11.8 | 4 |
| pEA808_5_7_QNR | 76.2 | 3.4 | 4 | pEA864_173_175_KLE | 20.3 | 2.8 | 2 |
| pEA809_8_10_KTG | 111.0 | 7.1 | 4 | pEA865_176_178_NTL | 4.0 | 2.6 | 4 |
| pEA810_11_13_NSK | 95.5 | 27.3 | 2 | pEA866_179_181_QDI | 1.6 | 1.9 | 4 |
| pEA811_14_16_TQS | 85.9 | 14.0 | 4 | pEA867_182_184_IQE | 6.6 | 9.8 | 4 |
| pEA812_17_19_ASP | 82.0 | 29.4 | 4 | pEA868_185_187_NFP | 5.8 | 9.1 | 4 |
| pEA813_20_22_PPK | 83.2 | 16.2 | 4 | pEA869_188_190_NLA | 51.7 | 12.3 | 2 |
| pEA814_23_25_ERS | 70.2 | 17.8 | 4 | pEA870_191_193_RQA | 101.2 | 22.8 | 4 |
| pEA815_26_28_SSP | 73.1 | 13.3 | 4 | pEA871_194_196_NVQ | 92.7 | 9.9 | 4 |
| pEA816_29_31_ATE | 100.4 | 24.6 | 4 | pEA872_197_199_IQE | 30.0 | 5.4 | 2 |
| pEA817_32_34_QSW | 81.4 | 17.3 | 4 | pEA873_200_202_IQR | 2.5 | 3.3 | 4 |
| pEA818_35_37_MEN | 93.1 | 14.3 | 6 | pEA874_203_205_TPQ | 5.6 | 1.8 | 4 |
| pEA819_38_40_DFD | 95.9 | 14.3 | 6 | pEA875_206_208_RYS | 6.9 | 6.0 | 4 |
| pEA820_41_43_ELR | 97.3 | 13.3 | 6 | pEA876_209_211_SRR | 9.2 | 4.3 | 4 |
| pEA821_44_46_EEG | 58.5 | 18.6 | 4 | pEA877_212_214_ATP | 10.9 | 4.9 | 4 |
| pEA822_47_49_FRR | 76.1 | 4.5 | 2 | pEA878_215_217_RHI | 2.7 | 3.5 | 4 |
| pEA823_50_52_SNY | 53.8 | 19.4 | 2 | pEA879_218_220_IVR | 5.3 | 1.2 | 2 |
| pEA824_53_55_SEL | 9.0 | 10.3 | 4 | pEA880_221_223_FTK | 11.5 | 2.8 | 2 |
| pEA825_56_58_RED | 72.5 | 8.0 | 2 | pEA881_224_226_VEM | 53.4 | 17.6 | 4 |
| pEA826_59_61_IQT | 10.2 | 4.9 | 4 | pEA882_227_229_KEK | 49.5 | 14.6 | 2 |
| pEA827_62_64_KGK | 78.0 | 9.3 | 2 | pEA883_230_232_MLR | 4.3 | 6.2 | 4 |
| pEA828_65_67_EVE | 8.4 | 4.5 | 4 | pEA884_233_235_AAR | 36.6 | 13.4 | 4 |
| pEA829_68_70_NFE | 12.9 | 10.9 | 4 | pEA885_236_238_EKG | 76.8 | 8.4 | 4 |
| pEA830_71_73_KNL | 25.1 | 9.9 | 4 | pEA886_239_241_RVT | 4.1 | 2.8 | 4 |
| pEA831_74_76_EEC | 68.9 | 11.3 | 4 | pEA887_242_244_LKG | 4.2 | 4.4 | 4 |
| pEA832_77_79_ITR | 8.8 | 4.7 | 2 | pEA888_245_247_KPI | 3.9 | 3.5 | 4 |
| pEA833_80_82_ITN | 42.2 | 13.1 | 3 | pEA889_248_250_RLT | 5.2 | 8.0 | 4 |
| pEA834_83_85_TEK | 8.3 | 7.2 | 3 | pEA890_251_253_ADL | 3.4 | 3.4 | 4 |
| pEA835_86_88_CLK | 9.8 | 6.0 | 4 | pEA891_254_256_SAE | 68.8 | 11.0 | 4 |
| pEA836_89_91_ELM | 4.2 | 1.3 | 4 | pEA892_257_259_TLQ | 5.6 | 3.1 | 4 |
| pEA837_92_94_ELK | 0.7 | 1.0 | 2 | pEA893_260_262_ARR | 3.2 | 3.0 | 4 |
| pEA838_95_97_TKA | 103.1 | 10.5 | 4 | pEA894_263_265_EWG | 4.0 | 1.6 | 4 |
| pEA839_98_100_REL | 5.8 | 3.0 | 4 | pEA895_266_268_PIF | 2.5 | 2.5 | 4 |
| pEA840_101_103_REE | 33.0 | 7.7 | 4 | pEA896_269_271_NIL | 2.5 | 1.9 | 4 |
| pEA841_104_106_CRS | 7.7 | 0.5 | 4 | pEA897_272_274_KEK | 57.6 | 11.2 | 4 |
| pEA842_107_109_LRS | 9.4 | 4.6 | 4 | pEA898_275_277_NFQ | 13.2 | 1.3 | 4 |
| pEA843_110_112_RCD | 10.1 | 5.1 | 4 | pEA899_278_280_PRI | 3.4 | 3.5 | 4 |
| pEA844_113_115_QLE | 2.9 | 2.1 | 4 | pEA900_281_283_SYP | 5.9 | 3.9 | 4 |
| pEA845_116_118_ERV | 13.1 | 11.2 | 4 | pEA901_284_286_AKL | 2.8 | 2.6 | 4 |
| pEA846_119_121_SAM | 36.4 | 2.8 | 4 | pEA902_287_289_SFI | 4.0 | 5.3 | 4 |
| pEA847_122_124_EDE | 4.0 | 1.8 | 4 | pEA903_290_292_SEG | 3.0 | 2.8 | 4 |
| pEA848_125_127_MNE | 60.0 | 3.1 | 4 | pEA904_293_295_EIK | 63.0 | 20.9 | 4 |
| pEA849_128_130_MKR | 34.8 | 6.4 | 4 | pEA905_296_298_YFI | 2.0 | 1.7 | 4 |
| pEA850_131_133_EGK | 91.4 | 16.4 | 4 | pEA906_299_301_DKQ | 1.9 | 1.9 | 4 |
| pEA851_134_136_FRE | 86.5 | 10.5 | 4 | pEA907_302_304_MLR | 24.5 | 8.4 | 4 |
| pEA852_137_139_KRI | 50.8 | 5.2 | 4 | pEA908_305_307_DFV | 4.7 | 1.1 | 2 |
| pEA853_140_142_KRN | 35.5 | 7.8 | 4 | pEA909_308_310_TTR | 60.6 | 13.2 | 4 |
| pEA854_143_145_EQS | 63.2 | 11.4 | 4 | pEA910_311_313_PAL | 2.4 | 2.8 | 4 |
| pEA855_146_148_LQE | 13.9 | 5.4 | 3 | pEA911_314_316_KEL | 79.3 | 17.4 | 2 |
| pEA856_149_151_IWD | 5.0 | 6.2 | 4 | pEA912_317_319_LKE | 13.4 | 3.1 | 4 |
| pEA857_152_154_YVK | 3.9 | 2.2 | 4 | pEA913_320_322_ALN | 33.5 | 15.0 | 4 |
| pEA858_155_157_RPN | 7.1 | 5.9 | 4 | pEA914_323_325_MER | 37.9 | 9.6 | 4 |
| pEA859_158_160_LRL | 5.9 | 4.9 | 4 | pEA915_326_328_NNR | 76.5 | 22.1 | 4 |
| pEA860_161_163_IGV | 11.0 | 9.6 | 3 | pEA916_329_331_YQP | 107.3 | 14.5 | 6 |
| pEA861_164_166_PES | 11.4 | 1.3 | 2 | pEA917_332_334_LQN | 84.6 | 20.1 | 4 |
| pEA862_167_169_DVE | 45.8 | 0.3 | 2 | pEA918_335_337_HAK | 85.1 | 17.0 | 4 |
| | | | | pEA1361_338_M | 77.4 | 12.2 | 4 |

| ORF2 | | | | | | | |
|---|---|---|---|---|---|---|---|
| Mutant ID | retroT average | standard deviation | number of measure-ments | Mutant ID | retroT average | standard deviation | number of measure-ments |
| pEA1362_2_3_TG | 24.8 | 1.5 | 4 | pEA981_187_189_KST | 87.9 | 11.6 | 4 |
| pEA920_4_6_STS | 72.0 | 5.5 | 2 | pEA982_190_192_EYT | 3.9 | 2.8 | 4 |
| pEA921_7_9_HIT | 17.6 | 20.4 | 4 | pEA983_193_195_FFS | 2.4 | 2.2 | 4 |
| pEA922_10_12_ILT | 4.3 | 5.0 | 4 | pEA984_196_198_APH | 10.3 | 5.2 | 4 |
| pEA923_13_15_LNI | 6.3 | 4.3 | 4 | pEA985_199_201_HTY | 6.0 | 1.1 | 2 |
| pEA924_16_18_NGL | 4.8 | 3.0 | 4 | pEA986_202_204_SKI | 7.5 | 5.1 | 4 |
| pEA925_19_21_NSA | 8.6 | 3.4 | 4 | pEA987_205_207_DHI | 1.9 | 1.8 | 4 |
| pEA926_22_24_IKR | 10.7 | 12.9 | 4 | pEA988_208_210_VGS | 6.6 | 5.1 | 4 |
| pEA927_25_27_HRL | 36.5 | 8.7 | 3 | pEA989_211_213_KAL | 6.0 | 3.4 | 4 |
| pEA928_28_30_ASW | 18.3 | 2.6 | 4 | pEA990_214_216_LSK | 61.5 | 23.9 | 4 |
| pEA929_31_33_IKS | 10.0 | 11.6 | 4 | pEA991_217_219_CKR | 91.6 | 29.0 | 4 |
| pEA930_34_36_QDP | 42.7 | 6.0 | 2 | pEA992_220_222_TEI | 45.9 | 8.8 | 4 |
| pEA931_37_39_SVC | 0.0 | 0.0 | 2 | pEA993_223_225_ITN | 33.4 | 8.6 | 4 |
| pEA932_40_42_CIQ | 2.3 | 2.5 | 4 | pEA994_226_228_YLS | 16.8 | 5.1 | 2 |
| pEA933_43_45_ETH | 1.7 | 2.1 | 4 | pEA995_229_231_DHS | 2.9 | 3.0 | 4 |
| pEA934_46_48_LTC | 5.8 | 3.9 | 4 | pEA996_232_234_AIK | 13.7 | 7.7 | 4 |
| pEA935_49_51_RDT | 81.1 | 3.7 | 4 | pEA997_235_237_LEL | 7.3 | 1.8 | 2 |
| pEA936_52_54_HRL | 4.2 | 5.0 | 4 | pEA998_238_240_RIK | 91.3 | 28.2 | 4 |
| pEA937_55_57_KIK | 15.0 | 7.0 | 4 | pEA999_241_243_NLT | 103.9 | 28.5 | 6 |
| pEA938_58_60_GWR | 3.9 | 0.8 | 2 | pEA1000_244_246_QSR | 60.4 | 13.1 | 2 |
| pEA939_61_63_KIY | 0.0 | 0.0 | 2 | pEA1001_247_249_STT | 87.9 | 26.3 | 4 |
| pEA940_64_66_QAN | 83.8 | 20.3 | 4 | pEA1002_250_252_WKL | 4.5 | 4.8 | 4 |
| pEA941_67_69_GKQ | 52.0 | 11.5 | 4 | pEA1003_253_255_NNL | 41.7 | 9.8 | 4 |
| pEA942_70_72_KKA | 7.2 | 4.6 | 4 | pEA1004_256_258_LLN | 9.6 | 2.0 | 4 |
| pEA943_73_75_GVA | 3.5 | 3.5 | 4 | pEA1005_259_261_DYW | 29.4 | 15.9 | 4 |
| pEA944_76_78_ILV | 1.6 | 1.8 | 4 | pEA1006_262_264_VHN | 106.1 | 12.5 | 2 |
| pEA945_79_81_SDK | 2.2 | 1.3 | 4 | pEA1007_265_267_EMK | 21.5 | 7.7 | 4 |
| pEA946_82_84_TDF | 6.3 | 0.1 | 2 | pEA1008_268_270_AEI | 13.0 | 9.9 | 3 |
| pEA947_85_87_KPT | 17.8 | 20.7 | 4 | pEA1009_271_273_KMF | 7.3 | 6.3 | 4 |
| pEA948_88_90_KIK | 10.0 | 2.3 | 4 | pEA1010_274_276_FET | 16.2 | 2.7 | 2 |
| pEA949_91_93_RDK | 10.0 | 7.6 | 2 | pEA1011_277_279_NEN | 12.2 | 6.9 | 4 |
| pEA950_94_96_EGH | 3.1 | 2.0 | 4 | pEA1012_280_282_KDT | 27.6 | 6.4 | 4 |
| pEA951_97_99_YIM | 2.3 | 2.5 | 4 | pEA1013_283_285_TYQ | 18.6 | 11.7 | 4 |
| pEA952_100_102_VKG | 3.4 | 1.8 | 4 | pEA1014_286_288_NLW | 2.6 | 2.7 | 4 |
| pEA953_103_105_SIQ | 27.8 | 8.3 | 4 | pEA1015_289_291_DAF | 3.2 | 2.2 | 4 |
| pEA954_106_108_QEE | 32.9 | 0.6 | 2 | pEA1016_292_294_KAV | 2.5 | 2.8 | 4 |
| pEA955_109_111_LTI | 5.3 | 4.6 | 3 | pEA1017_295_297_CRG | 4.8 | 4.2 | 4 |
| pEA956_112_114_LNI | 2.5 | 2.9 | 4 | pEA1018_298_300_KFI | 3.0 | 3.2 | 4 |
| pEA957_115_117_YAP | 2.2 | 1.5 | 4 | pEA1019_301_303_ALN | 103.0 | 25.1 | 4 |
| pEA958_118_120_NTG | 24.1 | 7.9 | 4 | pEA1020_304_306_AYK | 73.5 | 18.6 | 4 |
| pEA959_121_123_APR | 65.5 | 4.5 | 2 | pEA1021_307_309_RKQ | 62.5 | 4.2 | 2 |
| pEA960_124_126_FIK | 4.8 | 3.6 | 4 | pEA1022_310_312_ERS | 64.3 | 16.7 | 4 |
| pEA961_127_129_QVL | 2.1 | 2.1 | 4 | pEA1023_313_315_KID | 62.2 | 20.1 | 4 |
| pEA962_130_132_SDL | 39.8 | 3.6 | 2 | pEA1024_316_318_TLT | 59.4 | 5.4 | 4 |
| pEA963_133_135_QRD | 23.4 | 27.0 | 4 | pEA1025_319_321_SQL | 72.9 | 13.9 | 3 |
| pEA964_136_138_LDS | 7.2 | 0.1 | 2 | pEA1026_322_324_KEL | 68.7 | 14.0 | 4 |
| pEA965_139_141_HTL | 2.7 | 3.1 | 4 | pEA1027_325_327_EKQ | 86.9 | 7.9 | 4 |
| pEA966_142_144_IMG | 2.4 | 2.4 | 4 | pEA1028_328_330_EQT | 76.7 | 18.9 | 4 |
| pEA967_145_147_DFN | 4.0 | 4.6 | 4 | pEA1029_331_333_HSK | 82.8 | 20.4 | 4 |
| pEA968_148_150_TPL | 6.2 | 3.5 | 4 | pEA1030_334_336_ASR | 93.7 | 23.7 | 4 |
| pEA969_151_153_STL | 63.8 | 21.4 | 4 | pEA1031_337_339_RQE | 19.6 | 9.4 | 4 |
| pEA970_154_156_DRS | 2.5 | 2.9 | 4 | pEA1032_340_342_ITK | 31.1 | 3.5 | 4 |
| pEA971_157_159_TRQ | 51.5 | 15.9 | 4 | pEA1033_343_345_IRA | 29.6 | 8.2 | 4 |
| pEA972_160_162_KVN | 48.1 | 16.4 | 4 | pEA1034_346_348_ELK | 14.2 | 9.9 | 4 |
| pEA973_163_165_KDT | 96.8 | 11.9 | 4 | pEA1035_349_351_EIE | 59.0 | 8.5 | 2 |
| pEA974_166_168_QEL | 19.1 | 3.1 | 4 | pEA1036_352_354_TQK | 53.5 | 8.1 | 4 |
| pEA975_169_171_NSA | 79.4 | 21.8 | 4 | pEA1037_355_357_TLQ | 30.7 | 2.8 | 4 |
| pEA976_172_174_LHQ | 25.0 | 14.2 | 4 | pEA1038_358_360_KIN | 56.8 | 31.2 | 4 |
| pEA977_175_177_ADL | 8.4 | 0.7 | 2 | pEA1039_361_363_ESR | 31.5 | 15.2 | 3 |
| pEA978_178_180_IDI | 2.8 | 3.2 | 4 | pEA1040_364_366_SWF | 11.7 | 1.9 | 3 |
| pEA979_181_183_YRT | 2.9 | 2.1 | 4 | pEA1041_367_369_FER | 23.6 | 1.3 | 2 |
| pEA980_184_186_LHP | 4.9 | 0.7 | 2 | pEA1042_370_372_INK | 17.3 | 7.1 | 4 |

**ORF2**

| Mutant ID | retroT average | standard deviation | number of measure-ments | Mutant ID | retroT average | standard deviation | number of measure-ments |
|---|---|---|---|---|---|---|---|
| pEA1043_373_375_IDR | 61.2 | 11.0 | 4 | pEA1101_547_549_LAN | 53.5 | 3.8 | 4 |
| pEA1044_376_378_PLA | 41.7 | 13.5 | 4 | pEA1102_550_552_RIQ | 8.1 | 2.3 | 4 |
| pEA1045_379_381_RLI | 65.1 | 8.6 | 4 | pEA1103_553_555_QHI | 88.6 | 17.3 | 4 |
| pEA1046_382_384_KKK | 56.9 | 0.4 | 2 | pEA1104_556_558_KKL | 43.4 | 4.8 | 4 |
| pEA1047_385_387_REK | 58.6 | 1.3 | 2 | pEA1105_559_561_IHH | 16.3 | 11.5 | 4 |
| pEA1048_388_390_NQI | 59.8 | 20.4 | 4 | pEA1106_562_564_DQV | 6.2 | 3.5 | 4 |
| pEA1049_391_393_DTI | 60.3 | 12.5 | 6 | pEA1107_565_567_GFI | 20.4 | 7.8 | 4 |
| pEA1050_394_396_KND | 76.5 | 33.4 | 4 | pEA1108_568_570_PGM | 53.3 | 9.2 | 4 |
| pEA1051_397_399_KGD | 55.1 | 24.2 | 4 | pEA1109_571_573_QGW | 15.8 | 3.1 | 4 |
| pEA1052_400_402_ITT | 22.5 | 7.0 | 4 | pEA1110_574_576_FNI | 36.0 | 12.5 | 4 |
| pEA1053_403_405_DPT | 45.1 | 7.8 | 4 | pEA1111_577_579_RKS | 4.7 | 1.1 | 4 |
| pEA1054_406_408_EIQ | 14.4 | 5.0 | 4 | pEA1112_580_582_INV | 30.0 | 8.2 | 4 |
| pEA1055_409_411_TTI | 27.7 | 1.0 | 2 | pEA1113_583_585_IQH | 61.4 | 4.7 | 4 |
| pEA1056_412_414_REY | 27.9 | 3.8 | 3 | pEA1114_586_588_INR | 57.4 | 17.8 | 4 |
| pEA1057_415_417_YKH | 33.2 | 6.6 | 4 | pEA1115_589_591_AKD | 78.6 | 11.3 | 4 |
| pEA1058_418_420_LYA | 30.7 | 6.0 | 4 | pEA1116_592_594_KNH | 53.5 | 8.9 | 4 |
| pEA1059_421_423_NKL | 65.6 | 10.7 | 4 | pEA1117_595_597_MII | 6.3 | 2.0 | 4 |
| pEA1060_424_426_ENL | 49.6 | 7.7 | 4 | pEA1118_598_600_SID | 5.3 | 1.3 | 4 |
| pEA1061_427_429_EEM | 38.0 | 9.4 | 4 | pEA1119_601_603_AEK | 62.1 | 9.5 | 4 |
| pEA1062_430_432_DTF | 20.1 | 20.2 | 4 | pEA1120_604_606_AFD | 3.8 | 1.5 | 4 |
| pEA1063_433_435_LDT | 5.5 | 4.4 | 4 | pEA1121_607_609_KIQ | 38.7 | 6.4 | 4 |
| pEA1064_436_438_YTL | 18.5 | 14.5 | 4 | pEA1122_610_612_QPF | 5.1 | 0.6 | 2 |
| pEA1065_439_441_PRL | 22.6 | 34.5 | 4 | pEA1123_613_615_MLK | 6.4 | 1.9 | 4 |
| pEA1066_442_444_NQE | 73.3 | 11.8 | 4 | pEA1124_616_618_TLN | 35.5 | 3.2 | 4 |
| pEA1067_445_447_EVE | 42.9 | 16.2 | 4 | pEA1125_619_621_KLG | 13.0 | 2.1 | 3 |
| pEA1068_448_450_SLN | 30.3 | 12.9 | 3 | pEA1126_622_624_IDG | 7.9 | 4.3 | 4 |
| pEA1069_451_453_RPI | 30.8 | 33.2 | 4 | pEA1127_625_627_TYF | 18.8 | 7.9 | 4 |
| pEA1070_454_456_TGS | 87.5 | 18.3 | 4 | pEA1128_628_630_KII | 4.3 | 1.4 | 4 |
| pEA1071_457_459_EIV | 3.4 | 2.5 | 4 | pEA1129_631_633_RAI | 25.4 | 6.2 | 4 |
| pEA1072_460_462_AII | 16.3 | 4.1 | 3 | pEA1130_634_636_YDK | 13.6 | 1.0 | 4 |
| pEA1073_463_465_NSL | 12.6 | 12.9 | 3 | pEA1131_637_639_PTA | 62.5 | 12.6 | 4 |
| pEA1074_466_468_PTK | 40.0 | 4.9 | 2 | pEA1132_640_642_NII | 4.1 | 0.8 | 4 |
| pEA1075_469_471_KSP | 61.9 | 13.8 | 2 | pEA1133_643_645_LNG | 13.0 | 2.3 | 4 |
| pEA1076_472_474_GPD | 3.5 | 2.2 | 4 | pEA1134_646_648_QKL | 44.5 | 4.6 | 4 |
| pEA1077_475_477_GFT | 7.4 | 6.9 | 4 | pEA1135_649_651_EAF | 6.2 | 3.5 | 4 |
| pEA1078_478_480_AEF | 18.5 | 10.6 | 4 | pEA1136_652_654_PLK | 35.3 | 5.1 | 4 |
| pEA1079_481_483_YQR | 14.4 | 13.8 | 4 | pEA1137_655_657_TGT | 10.7 | 1.1 | 4 |
| pEA1080_484_486_YKE | 8.0 | 2.4 | 2 | pEA1138_658_660_RQG | 8.7 | 5.6 | 4 |
| pEA1081_487_489_ELV | 9.7 | 6.4 | 4 | pEA1139_661_663_CPL | 5.6 | 1.5 | 4 |
| pEA1082_490_492_PFL | 8.1 | 7.5 | 4 | pEA1140_664_666_SPL | 4.8 | 1.3 | 4 |
| pEA1083_493_495_LKL | 6.2 | 2.0 | 4 | pEA1141_667_669_LFN | 11.6 | 3.0 | 4 |
| pEA1084_496_498_FQS | 28.1 | 8.4 | 4 | pEA1142_670_672_IVL | 13.9 | 5.4 | 4 |
| pEA1085_499_501_IEK | 79.3 | 27.2 | 4 | pEA1143_673_675_EVL | 4.9 | 1.2 | 4 |
| pEA1086_502_504_EGI | 38.1 | 9.5 | 4 | pEA1144_676_678_ARA | 123.2 | 15.7 | 4 |
| pEA1087_505_507_LPN | 8.2 | 1.6 | 4 | pEA1145_679_681_IRQ | 5.9 | 1.0 | 4 |
| pEA1088_508_510_SFY | 8.9 | 2.4 | 4 | pEA1146_682_684_EKE | 29.2 | 5.8 | 4 |
| pEA1089_511_513_EAS | 41.1 | 6.5 | 4 | pEA1147_685_687_IKG | 4.4 | 1.4 | 4 |
| pEA1090_514_516_IIL | 5.9 | 2.3 | 4 | pEA1148_688_690_IQL | 9.1 | 1.9 | 4 |
| pEA1091_517_519_IPK | 8.2 | 0.9 | 3 | pEA1149_691_693_GKE | 51.7 | 12.3 | 4 |
| pEA1092_520_522_PGR | 9.9 | 6.1 | 4 | pEA1150_694_696_EVK | 3.4 | 1.1 | 4 |
| pEA1093_523_525_DTT | 80.1 | 24.7 | 4 | pEA1151_697_699_LSL | 36.9 | 4.3 | 4 |
| pEA1094_526_528_KKE | 82.6 | 6.7 | 4 | pEA1152_700_702_FAD | 9.4 | 6.3 | 4 |
| pEA1095_529_531_NFR | 5.6 | 0.7 | 4 | pEA1153_703_705_DMI | 4.0 | 1.3 | 4 |
| pEA1096_532_534_PIS | 5.7 | 2.6 | 4 | pEA1154_706_708_VYL | 7.3 | 3.0 | 3 |
| pEA1097_535_537_LMN | 9.7 | 5.4 | 2 | pEA1155_709_711_ENP | 15.8 | 7.2 | 3 |
| pEA1098_538_540_IDA | 8.0 | 6.7 | 4 | pEA1156_712_714_IVS | 34.3 | 4.1 | 4 |
| pEA1099_541_543_KIL | 4.5 | 1.5 | 4 | pEA1157_715_717_AQN | 88.0 | 12.9 | 3 |
| pEA1100_544_546_NKI | 58.1 | 7.5 | 4 | pEA1158_718_720_LLK | 7.3 | 4.0 | 4 |

| ORF2 | | | | | | | |
|---|---|---|---|---|---|---|---|
| Mutant ID | retroT average | standard deviation | number of measure-ments | Mutant ID | retroT average | standard deviation | number of measure-ments |
| pEA1159_721_723_LIS | 6.0 | 3.1 | 4 | pEA1221_907_909_IMP | 112.1 | 4.0 | 4 |
| pEA1160_724_726_NFS | 5.4 | 1.1 | 4 | pEA1222_910_912_HIY | 88.5 | 4.0 | 4 |
| pEA1161_727_729_KVS | 20.1 | 7.2 | 4 | pEA1223_913_915_NYL | 105.1 | 4.0 | 4 |
| pEA1162_730_732_GYK | 5.6 | 2.6 | 4 | pEA1224_916_918_IFD | 7.3 | 4.0 | 4 |
| pEA1163_733_735_INV | 11.4 | 3.2 | 4 | pEA1225_919_921_KPE | 95.3 | 4.0 | 4 |
| pEA1164_736_738_QKS | 5.3 | 1.9 | 4 | pEA1226_922_924_KNK | 79.0 | 4.0 | 4 |
| pEA1165_739_741_QAF | 8.5 | 2.9 | 4 | pEA1227_925_927_QWG | 112.0 | 4.0 | 4 |
| pEA1166_742_744_LYT | 6.4 | 2.2 | 4 | pEA1228_928_930_KDS | 73.2 | 4.0 | 4 |
| pEA1167_745_747_NNR | 59.7 | 10.0 | 4 | pEA1229_931_933_LFN | 48.9 | 4.0 | 4 |
| pEA1168_748_750_QTE | 92.1 | 7.4 | 4 | pEA1230_934_936_KWC | 75.3 | 4.0 | 4 |
| pEA1169_751_753_SQI | 119.7 | 4.0 | 4 | pEA1231_937_939_WEN | 64.5 | 4.0 | 4 |
| pEA1170_754_756_MGE | 115.8 | 11.8 | 6 | pEA1232_940_942_WLA | 18.7 | 4.0 | 4 |
| pEA1171_757_759_LPF | 22.2 | 0.9 | 4 | pEA1233_943_945_ICR | 96.0 | 4.0 | 4 |
| pEA1172_760_762_TIA | 92.5 | 27.1 | 4 | pEA1234_946_948_KLK | 97.5 | 4.0 | 4 |
| pEA1173_763_765_SKR | 66.0 | 14.7 | 4 | pEA1235_949_951_LDP | 70.9 | 4.0 | 4 |
| pEA1174_766_768_IKY | 10.2 | 3.2 | 4 | pEA1236_952_954_FLT | 16.2 | 4.0 | 4 |
| pEA1175_769_771_LGI | 4.3 | 2.4 | 4 | pEA1237_955_957_PYT | 3.8 | 4.0 | 4 |
| pEA1176_772_774_QLT | 20.0 | 3.0 | 4 | pEA1238_958_960_KIN | 7.9 | 4.0 | 4 |
| pEA1177_775_777_RDV | 78.3 | 17.3 | 4 | pEA1239_961_963_SRW | 9.7 | 4.0 | 4 |
| pEA1178_778_780_KDL | 44.3 | 11.3 | 4 | pEA1240_964_966_IKD | 42.0 | 4.0 | 4 |
| pEA1179_781_783_FKE | 82.7 | 8.9 | 4 | pEA1241_967_969_LNV | 7.6 | 4.0 | 4 |
| pEA1180_784_786_NYK | 14.2 | 2.7 | 4 | pEA1242_970_972_KPK | 64.2 | 4.0 | 4 |
| pEA1181_787_789_PLL | 42.3 | 2.2 | 4 | pEA1243_973_975_TIK | 13.8 | 2.0 | 2 |
| pEA1182_790_792_KEI | 53.8 | 5.4 | 4 | pEA1244_976_978_TLE | 27.6 | 4.0 | 4 |
| pEA1183_793_795_KEE | 75.4 | 12.4 | 4 | pEA1245_979_981_ENL | 63.0 | 4.0 | 4 |
| pEA1184_796_798_TNK | 104.1 | 15.3 | 4 | pEA1246_982_984_GIT | 25.0 | 3.0 | 3 |
| pEA1185_799_801_WKN | 62.1 | 6.6 | 4 | pEA1247_985_987_IQD | 50.7 | 4.0 | 4 |
| pEA1186_802_804_IPC | 79.9 | 9.3 | 4 | pEA1248_988_990_IGV | 10.2 | 4.0 | 4 |
| pEA1187_805_807_SWV | 24.5 | 9.2 | 4 | pEA1249_991_993_GKD | 97.4 | 4.0 | 4 |
| pEA1188_808_810_GRI | 34.3 | 4.9 | 4 | pEA1250_994_996_FMS | 8.0 | 4.0 | 4 |
| pEA1189_811_813_NIV | 41.4 | 7.7 | 4 | pEA1251_997_999_KTP | 93.4 | 4.0 | 4 |
| pEA1190_814_816_KMA | 35.3 | 15.8 | 4 | pEA1252_1000_1002_KAM | 103.2 | 4.0 | 4 |
| pEA1191_817_819_ILP | 10.7 | 3.3 | 4 | pEA1253_1003_1005_ATK | 108.8 | 4.0 | 4 |
| pEA1192_820_822_KVI | 19.4 | 3.8 | 4 | pEA1254_1006_1008_DKI | 77.3 | 4.0 | 4 |
| pEA1193_823_825_YRF | 100.1 | 26.0 | 4 | pEA1255_1009_1011_DKW | 11.0 | 4.0 | 4 |
| pEA1194_826_828_NAI | 55.8 | 3.9 | 4 | pEA1256_1012_1014_DLI | 10.6 | 4.0 | 4 |
| pEA1195_829_831_PIK | 5.2 | 0.4 | 4 | pEA1257_1015_1017_KLK | 10.7 | 4.0 | 4 |
| pEA1196_832_834_LPM | 66.1 | 21.2 | 4 | pEA1258_1018_1020_SFC | 6.4 | 4.0 | 4 |
| pEA1197_835_837_TFF | 34.8 | 17.9 | 4 | pEA1259_1021_1023_TAK | 92.8 | 4.0 | 4 |
| pEA1198_838_840_TEL | 67.4 | 13.5 | 4 | pEA1260_1024_1026_ETT | 91.0 | 4.0 | 4 |
| pEA1199_841_843_EKT | 82.5 | 6.4 | 4 | pEA1261_1027_1029_IRV | 109.8 | 4.0 | 4 |
| pEA1200_844_846_TLK | 97.6 | 18.3 | 2 | pEA1262_1030_1032_NRQ | 71.9 | 3.0 | 3 |
| pEA1201_847_849_FIW | 8.2 | 2.4 | 4 | pEA1263_1033_1035_PTT | 125.7 | 4.0 | 4 |
| pEA1202_850_852_NQK | 83.3 | 14.9 | 4 | pEA1264_1036_1038_WEK | 28.5 | 4.0 | 4 |
| pEA1203_853_855_RAR | 18.9 | 4.0 | 4 | pEA1265_1039_1041_IFA | 7.8 | 4.0 | 4 |
| pEA1204_856_858_IAK | 52.0 | 4.0 | 4 | pEA1266_1042_1044_TYS | 106.5 | 4.0 | 4 |
| pEA1205_859_861_SIL | 56.0 | 4.0 | 4 | pEA1267_1045_1047_SDK | 40.6 | 4.0 | 4 |
| pEA1206_862_864_SQK | 72.9 | 4.0 | 4 | pEA1268_1048_1050_GLI | 14.6 | 4.0 | 4 |
| pEA1207_865_867_NKA | 122.1 | 4.0 | 4 | pEA1269_1051_1053_SRI | 55.3 | 2.0 | 2 |
| pEA1208_868_870_GGI | 60.4 | 4.0 | 4 | pEA1270_1054_1056_YNE | 66.0 | 4.0 | 4 |
| pEA1209_871_873_TLP | 69.2 | 4.0 | 4 | pEA1271_1057_1059_LKQ | 82.3 | 4.0 | 4 |
| pEA1210_874_876_DFK | 6.7 | 4.0 | 4 | pEA1272_1060_1062_IYK | 66.5 | 2.0 | 2 |
| pEA1211_877_879_LYY | 5.6 | 4.0 | 4 | pEA1273_1063_1065_KKT | 107.5 | 4.0 | 4 |
| pEA1212_880_882_KAT | 59.9 | 4.0 | 4 | pEA1274_1066_1068_NNP | 56.6 | 4.0 | 4 |
| pEA1213_883_885_VTK | 62.3 | 4.0 | 4 | pEA1275_1069_1071_IKK | 42.6 | 2.0 | 2 |
| pEA1214_886_888_TAW | 36.4 | 4.0 | 4 | pEA1276_1072_1074_WAK | 17.6 | 4.0 | 4 |
| pEA1215_889_891_YWY | 3.9 | 4.0 | 4 | pEA1277_1075_1077_DMN | 5.4 | 4.0 | 4 |
| pEA1216_892_894_QNR | 37.8 | 4.0 | 4 | pEA1278_1078_1080_RHF | 14.2 | 4.0 | 4 |
| pEA1217_895_897_DID | 54.9 | 4.0 | 4 | pEA1279_1081_1083_SKE | 93.5 | 4.0 | 4 |
| pEA1218_898_900_QWN | 59.9 | 4.0 | 4 | pEA1280_1084_1086_DIY | 65.1 | 4.0 | 4 |
| pEA1219_901_903_RTE | 68.9 | 4.0 | 4 | pEA1281_1087_1089_AAK | 64.0 | 4.0 | 4 |
| pEA1220_904_906_PSE | 119.5 | 4.0 | 4 | pEA1282_1090_1092_KHM | 14.9 | 4.0 | 4 |

| ORF2 | | | | | | | |
|---|---|---|---|---|---|---|---|
| **Mutant ID** | **retroT average** | **standard deviation** | **number of measure-ments** | **Mutant ID** | **retroT average** | **standard deviation** | **number of measure-ments** |
| pEA1283_1093_1095_KKC | 65.4 | 4.0 | 4 | pEA1314_1186_1188_SCC | 97.2 | 5.0 | 5 |
| pEA1284_1096_1098_SSS | 43.6 | 4.0 | 4 | pEA1315_1189_1191_YKD | 94.0 | 4.0 | 4 |
| pEA1285_1099_1101_LAI | 50.2 | 4.0 | 4 | pEA1316_1192_1194_TCT | 63.9 | 4.0 | 4 |
| pEA1286_1102_1104_REM | 24.2 | 4.0 | 4 | pEA1317_1195_1197_RMF | 19.2 | 4.0 | 4 |
| pEA1287_1105_1107_QIK | 27.9 | 4.0 | 4 | pEA1318_1198_1200_IAA | 102.4 | 6.0 | 6 |
| pEA1288_1108_1110_TTM | 43.7 | 4.0 | 4 | pEA1319_1201_1203_LFT | 82.2 | 4.0 | 4 |
| pEA1289_1111_1113_RYH | 5.3 | 2.0 | 2 | pEA1320_1204_1206_IAK | 68.5 | 2.0 | 2 |
| pEA1290_1114_1116_LTP | 21.1 | 2.0 | 2 | pEA1321_1207_1209_TWN | 13.1 | 2.0 | 2 |
| pEA1291_1117_1119_VRM | 70.3 | 4.0 | 4 | pEA1322_1210_1212_QPK | 85.6 | 2.0 | 2 |
| pEA1292_1120_1122_AII | 116.5 | 6.0 | 6 | pEA1323_1213_1215_CPT | 63.0 | 2.0 | 2 |
| pEA1293_1123_1125_KKS | 92.7 | 4.0 | 4 | pEA1324_1216_1218_MID | 91.0 | 4.0 | 4 |
| pEA1294_1126_1128_GNN | 97.2 | 4.0 | 4 | pEA1325_1219_1221_WIK | 24.9 | 4.0 | 4 |
| pEA1295_1129_1131_RCW | 7.9 | 4.0 | 4 | pEA1326_1222_1224_KMW | 7.8 | 4.0 | 4 |
| pEA1296_1132_1134_RGC | 6.3 | 4.0 | 4 | pEA1327_1225_1227_HIY | 24.1 | 4.0 | 4 |
| pEA1297_1135_1137_GEI | 126.9 | 4.0 | 4 | pEA1328_1228_1230_TME | 4.6 | 4.0 | 4 |
| pEA1298_1138_1140_GTL | 82.0 | 4.0 | 4 | pEA1329_1231_1233_YYA | 23.3 | 2.0 | 2 |
| pEA1299_1141_1143_LHC | 5.2 | 4.0 | 4 | pEA1330_1234_1236_AIK | 50.8 | 2.0 | 2 |
| pEA1300_1144_1146_WWD | 18.7 | 4.0 | 4 | pEA1331_1237_1239_NDE | 61.1 | 3.0 | 3 |
| pEA1301_1147_1149_CKL | 11.3 | 4.0 | 4 | pEA1332_1240_1242_FIS | 59.6 | 4.0 | 4 |
| pEA1302_1150_1152_VQP | 46.3 | 4.0 | 4 | pEA1333_1243_1245_FVG | 28.4 | 3.0 | 3 |
| pEA1303_1153_1155_LWK | 10.9 | 4.0 | 4 | pEA1334_1246_1248_TWM | 16.3 | 4.0 | 4 |
| pEA1304_1156_1158_SVW | 23.6 | 4.0 | 4 | pEA1335_1249_1251_KLE | 20.6 | 2.0 | 2 |
| pEA1305_1159_1161_RFL | 87.2 | 4.0 | 4 | pEA1336_1252_1254_TII | 8.8 | 4.0 | 4 |
| pEA1306_1162_1164_RDL | 100.4 | 4.0 | 4 | pEA1337_1255_1257_LSK | 56.4 | 4.0 | 4 |
| pEA1307_1165_1167_ELE | 79.5 | 4.0 | 4 | pEA1338_1258_1260_LSQ | 82.5 | 3.0 | 3 |
| pEA1308_1168_1170_IPF | 49.7 | 4.0 | 4 | pEA1339_1261_1263_EQK | 104.1 | 6.0 | 6 |
| pEA1309_1171_1173_DPA | 69.7 | 4.0 | 4 | pEA1340_1264_1266_TKH | 101.8 | 8.0 | 8 |
| pEA1310_1174_1176_IPL | 21.3 | 4.0 | 4 | pEA1341_1267_1269_RIF | 48.8 | 4.0 | 4 |
| pEA1311_1177_1179_LGI | 4.5 | 4.0 | 4 | pEA1342_1270_1272_SLI | 80.8 | 4.0 | 4 |
| pEA1312_1180_1182_YPN | 86.7 | 4.0 | 4 | pEA1343_1273_1275_GGN | 58.4 | 8.0 | 8 |
| pEA1313_1183_1185_EYK | 109.3 | 4.0 | 4 | | | | |

**Supplemental Figure 3**

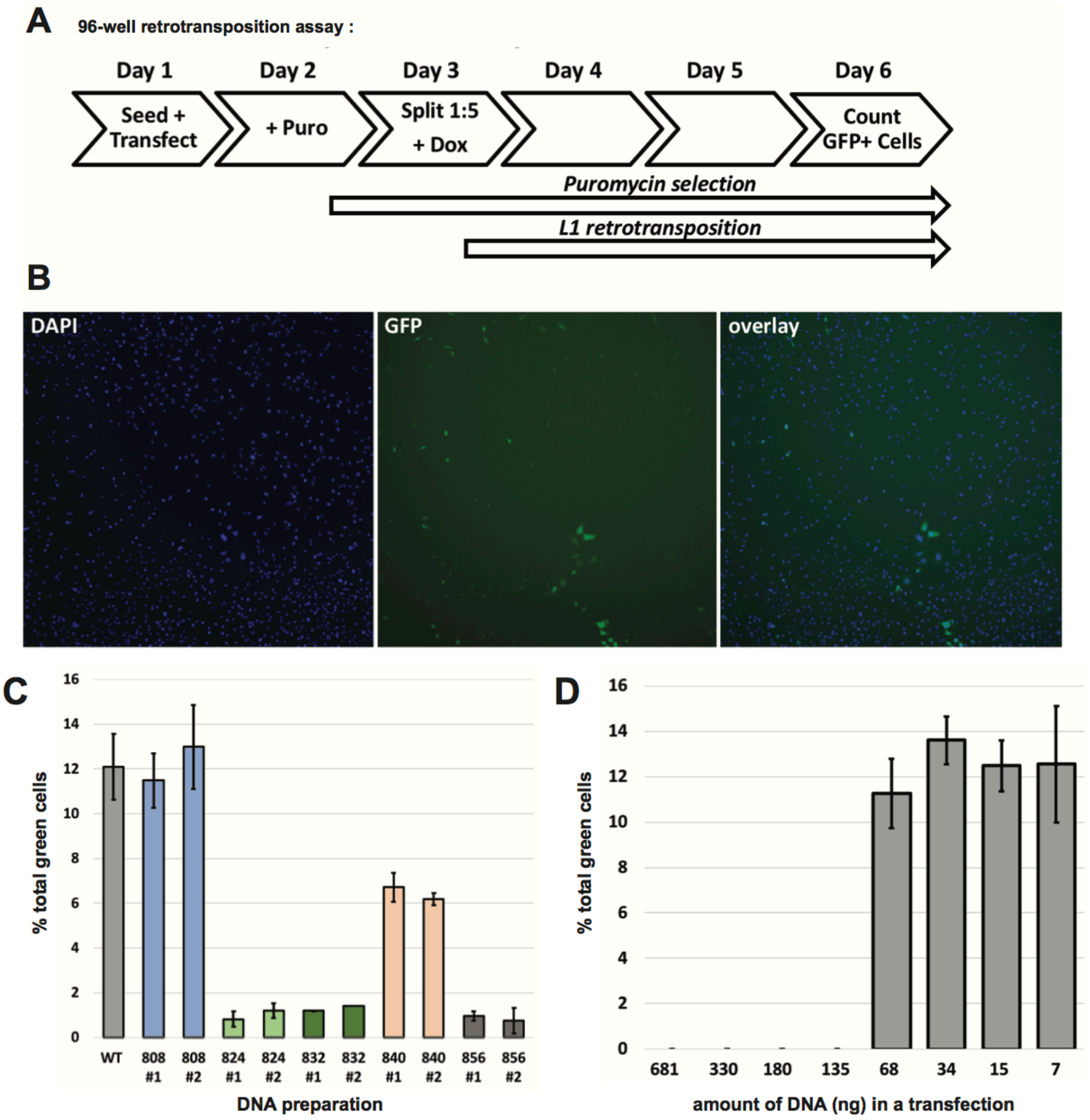

#### Supplemental Figure 4

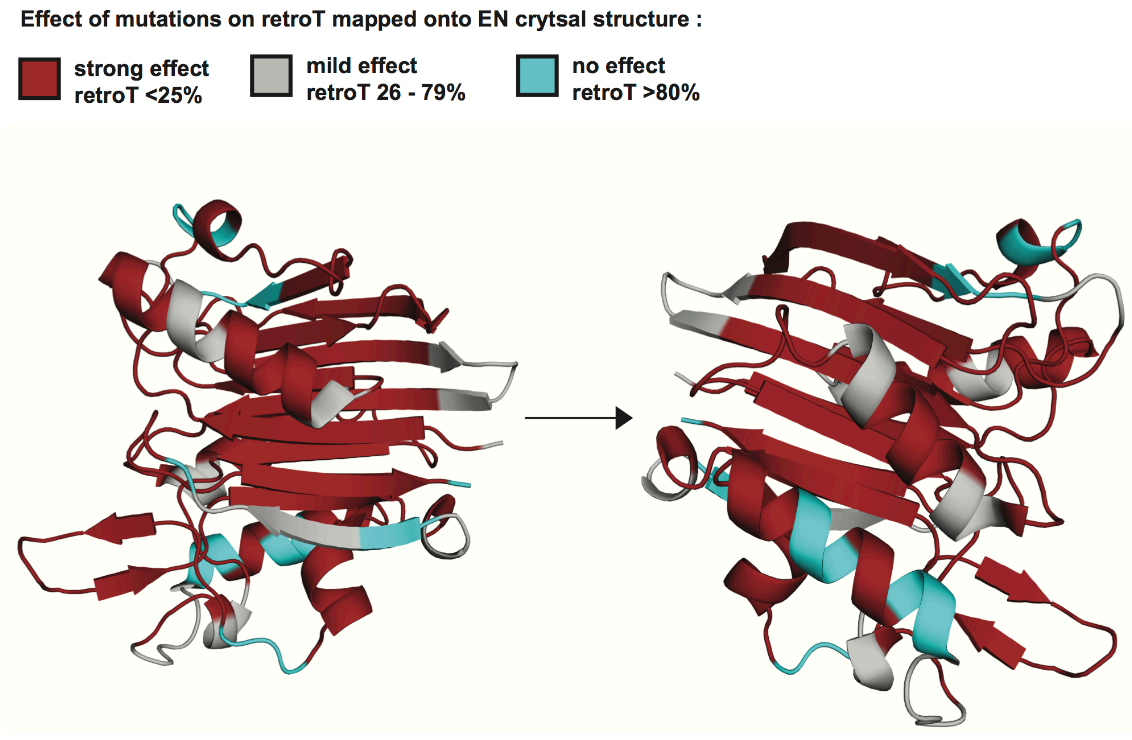

#### Supplemental Table 3

| ORF1 | | | |
|---|---|---|---|
| **Mutant ID** | **average** | **Mutant ID** | **average** |
| pEA807_2_4_GKK | 131.9 | pEA864_173_175_KLE | 49.6 |
| pEA808_5_7_QNR | 138.3 | pEA865_176_178_NTL | 22.2 |
| pEA809_8_10_KTG | 78.5 | pEA866_179_181_QDI | 3.2 |
| pEA810_11_13_NSK | 168.9 | pEA867_182_184_IQE | 8.7 |
| pEA811_14_16_TQS | 201.1 | pEA868_185_187_NFP | 11.5 |
| pEA812_17_19_ASP | 94.0 | pEA869_188_190_NLA | 133.5 |
| pEA813_20_22_PPK | 122.0 | pEA870_191_193_RQA | 120.4 |
| pEA814_23_25_ERS | 88.5 | pEA871_194_196_NVQ | 101.3 |
| pEA815_26_28_SSP | 94.9 | pEA872_197_199_IQE | 109.7 |
| pEA816_29_31_ATE | 93.3 | pEA873_200_202_IQR | 12.7 |
| pEA817_32_34_QSW | 151.2 | pEA874_203_205_TPQ | 90.0 |
| pEA818_35_37_MEN | *mutates epitope* | pEA875_206_208_RYS | 129.5 |
| pEA819_38_40_DFD | *mutates epitope* | pEA876_209_211_SRR | 108.2 |
| pEA820_41_43_ELR | *mutates epitope* | pEA877_212_214_ATP | 72.4 |
| pEA821_44_46_EEG | *mutates epitope* | pEA878_215_217_RHI | 24.1 |
| pEA822_47_49_FRR | 111.0 | pEA879_218_220_IVR | 4.9 |
| pEA823_50_52_SNY | 148.1 | pEA880_221_223_FTK | 46.7 |
| pEA824_53_55_SEL | 109.4 | pEA881_224_226_VEM | 106.4 |
| pEA825_56_58_RED | 102.9 | pEA882_227_229_KEK | 99.5 |
| pEA826_59_61_IQT | 110.6 | pEA883_230_232_MLR | 22.3 |
| pEA827_62_64_KGK | 153.9 | pEA884_233_235_AAR | 104.1 |
| pEA828_65_67_EVE | 104.4 | pEA885_236_238_EKG | 81.7 |
| pEA829_68_70_NFE | 139.7 | pEA886_239_241_RVT | 63.2 |
| pEA830_71_73_KNL | 161.1 | pEA887_242_244_LKG | 46.2 |
| pEA831_74_76_EEC | 93.5 | pEA888_245_247_KPI | 5.8 |
| pEA832_77_79_ITR | 196.1 | pEA889_248_250_RLT | 68.5 |
| pEA833_80_82_ITN | 160.9 | pEA890_251_253_ADL | 7.9 |
| pEA834_83_85_TEK | 106.3 | pEA891_254_256_SAE | 53.9 |
| pEA835_86_88_CLK | 187.7 | pEA892_257_259_TLQ | 109.0 |
| pEA836_89_91_ELM | 82.6 | pEA893_260_262_ARR | 81.2 |
| pEA837_92_94_ELK | 99.6 | pEA894_263_265_EWG | 10.6 |
| pEA838_95_97_TKA | 136.8 | pEA895_266_268_PIF | 13.4 |
| pEA839_98_100_REL | 119.7 | pEA896_269_271_NIL | 14.7 |
| pEA840_101_103_REE | 140.5 | pEA897_272_274_KEK | 118.9 |
| pEA841_104_106_CRS | 146.7 | pEA898_275_277_NFQ | 74.0 |
| pEA842_107_109_LRS | 175.6 | pEA899_278_280_PRI | 21.1 |
| pEA843_110_112_RCD | 188.7 | pEA900_281_283_SYP | 68.4 |
| pEA844_113_115_QLE | 147.2 | pEA901_284_286_AKL | 22.9 |
| pEA845_116_118_ERV | 90.0 | pEA902_287_289_SFI | 10.4 |
| pEA846_119_121_SAM | 87.4 | pEA903_290_292_SEG | 46.4 |
| pEA847_122_124_EDE | 169.1 | pEA904_293_295_EIK | 82.7 |
| pEA848_125_127_MNE | 134.7 | pEA905_296_298_YFI | 86.9 |
| pEA849_128_130_MKR | 175.3 | pEA906_299_301_DKQ | 19.4 |
| pEA850_131_133_EGK | 157.3 | pEA907_302_304_MLR | 89.1 |
| pEA851_134_136_FRE | 157.2 | pEA908_305_307_DFV | 47.8 |
| pEA852_137_139_KRI | 121.6 | pEA909_308_310_TTR | 126.1 |
| pEA853_140_142_KRN | 116.5 | pEA910_311_313_PAL | 42.4 |
| pEA854_143_145_EQS | 98.3 | pEA911_314_316_KEL | 89.8 |
| pEA855_146_148_LQE | 120.5 | pEA912_317_319_LKE | 39.6 |
| pEA856_149_151_IWD | 65.5 | pEA913_320_322_ALN | 119.0 |
| pEA857_152_154_YVK | 102.9 | pEA914_323_325_MER | 99.6 |
| pEA858_155_157_RPN | 93.2 | pEA915_326_328_NNR | 106.3 |
| pEA859_158_160_LRL | 11.4 | pEA916_329_331_YQP | 110.5 |
| pEA860_161_163_IGV | 12.6 | pEA917_332_334_LQN | 154.9 |
| pEA861_164_166_PES | 37.6 | pEA918_335_337_HAK | 91.9 |
| pEA862_167_169_DVE | 66.0 | pEA1361_338_M | 94.7 |
| pEA863_170_172_NGT | 104.0 | | |

#### Supplemental Table 4

| % of 3xala mutations that landed in each final functional category: | | | | |
|---|---|---|---|---|
| **ORF1p** | | | | |
| | **WT retroT high ORF1p** | **reduced retroT high ORF1p** | **no retroT high ORF1p** | **no retroT low ORF1p** |
| **Full Length Protein** | **16** | **31** | **29** | **24** |
| NTR | 9 | 4 | 0 | 0 |
| Coiled coil | 3 | 11 | 16 | 0 |
| RRM | 0 | 7 | 6 | 14 |
| CTD | 2 | 6 | 5 | 10 |

#### Supplemental Table 5

| | Mutants in each pool (1348 is WT) | | | | | | | | | | | | | | | |
|---|---|---|---|---|---|---|---|---|---|---|---|---|---|---|---|---|
| **Pool 1** | 807 | 815 | 823 | 831 | 839 | 847 | 855 | 863 | 869 | 877 | 885 | 893 | 901 | 909 | 917 | 1348 |
| **Pool 2** | 808 | 816 | 824 | 832 | 840 | 848 | 856 | 864 | 870 | 878 | 886 | 894 | 902 | 910 | 918 | 1348 |
| **Pool 3** | 809 | 817 | 825 | 833 | 841 | 849 | 857 | 871 | 879 | 887 | 895 | 903 | 911 | 1361 | | 1348 |
| **Pool 4** | 810 | 818 | 826 | 834 | 842 | 850 | 858 | 865 | 873 | 881 | 889 | 897 | 905 | 913 | | 1348 |
| **Pool 5** | 811 | 819 | 827 | 835 | 843 | 851 | 859 | 866 | 874 | 882 | 890 | 898 | 906 | 914 | | 1348 |
| **Pool 6** | 812 | 820 | 828 | 836 | 844 | 852 | 860 | 867 | 875 | 883 | 891 | 899 | 907 | 915 | | 1348 |
| **Pool 7** | 813 | 821 | 829 | 837 | 845 | 853 | 861 | 868 | 876 | 884 | 892 | 900 | 908 | 916 | | 1348 |
| **Pool 8** | 814 | 822 | 830 | 838 | 846 | 854 | 862 | 872 | 880 | 888 | 896 | 904 | 912 | | | 1348 |

#### Supplemental Table 6

| | ORF1p domain | retroT category | # of measurements | nucleolar phenotype (>5% positive) | aggregate blinded counting data | |
|---|---|---|---|---|---|---|
| | | | | | % nucleolar ORF1 positive cells | # of cells counted |
| **WT** | | | 4 | - | 0.82 | 122 |
| **pEA807_2_4_GKK** | | - | 2 | - | 0 | 41 |
| **pEA812_17_19_ASP** | **NTR** | ++ | 2 | - | 0 | 31 |
| **pEA814_23_25_ERS** | | + | 2 | - | 0 | 57 |
| **pEA816_29_31_ATE** | | ++ | 2 | - | 0 | 58 |
| **pEA824_53_55_SEL** | | - | 2 | - | 0 | 49 |
| **pEA826_59_61_IQT** | | - | 2 | - | 0 | 51 |
| **pEA828_65_67_EVE** | | - | 1 | - | 0 | 27 |
| **pEA829_68_70_NFE** | | - | 1 | - | 2.7 | 37 |
| **pEA832_77_79_ITR** | | - | 1 | - | 0 | 24 |
| **pEA833_80_82_ITN** | | + | 1 | - | 0 | 32 |
| **pEA834_83_85_TEK** | | - | 1 | - | 3.6 | 28 |
| **pEA835_86_88_CLK** | | - | 2 | + | 7.5 | 67 |
| **pEA836_89_91_ELM** | | - | 3 | - | 2.3 | 86 |
| **pEA837_92_94_ELK** | **CC** | - | 3 | - | 0 | 72 |
| **pEA838_95_97_TKA** | | ++ | 1 | - | 3.2 | 31 |
| **pEA839_98_100_REL** | | - | 1 | - | 0 | 30 |
| **pEA841_104_106_CRS** | | - | 1 | - | 0 | 21 |
| **pEA842_107_109_LRS** | | - | 1 | + | 24.1 | 29 |
| **pEA843_110_112_RCD** | | - | 1 | - | 0 | 21 |
| **pEA847_122_124_EDE** | | - | 1 | - | 0 | 39 |
| **pEA850_131_133_EGK** | | ++ | 1 | - | 0 | 22 |
| **pEA855_146_148_LQE** | | - | 1 | - | 0 | 27 |
| **pEA858_155_157_RPN** | | - | 1 | - | 0 | 37 |
| **pEA861_164_166_PES** | | - | 1 | - | 3.6 | 28 |
| **pEA866_179_181_QDI** | | - | 2 | - | 3.4 | 59 |
| **pEA870_191_193_RQA** | | ++ | 1 | - | 0 | 21 |
| **pEA874_203_205_TPQ** | | - | 1 | - | 0 | 33 |
| **pEA877_212_214_ATP** | **RRM** | - | 1 | - | 0 | 30 |
| **pEA880_221_223_FTK** | | - | 2 | - | 0 | 45 |
| **pEA886_239_241_RVT** | | - | 2 | - | 0 | 59 |
| **pEA887_242_244_LKG** | | - | 1 | - | 0 | 37 |
| **pEA889_248_250_RLT** | | - | 1 | - | 0 | 48 |
| **pEA893_260_262_ARR** | | - | 1 | - | 0 | 33 |
| **pEA898_275_277_NFQ** | | - | 1 | - | 0 | 37 |
| **pEA900_281_283_SYP** | | - | 2 | - | 3.8 | 52 |
| **pEA903_290_292_SEG** | **CTD** | - | 1 | - | 0 | 43 |
| **pEA908_305_307_DFV** | | - | 1 | - | 0 | 40 |
| **pEA910_311_313_PAL** | | - | 2 | - | 2.7 | 73 |
| **pEA911_314_316_KEL** | | + | 1 | - | 0 | 34 |
| **pEA914_323_325_MER** | | + | 1 | - | 0 | 26 |

**Supplemental Table 7**

| MAMMALIAN | NON-MAMMALIAN VERTEBRATE | | |
|---|---|---|---|
| HUMAN L1 - RP | LIZARD L1_AC1 | FROG L1-15 | ZEBRAFISH L1-7B |
| ARMADILLO | LIZARD L1_AC3B | FROG L1-17 | ZEBRAFISH L1-7C |
| COW | LIZARD L1_AC7 | FROG L1-18 | ZEBRAFISH L1-8 |
| DOG | LIZARD L1_AC8 | FROG L1-29 | ZEBRAFISH L1-10B |
| PIG | LIZARD L1_AC9 | FROG L1-32 | ZEBRAFISH L1-11A |
| ELEPHANT | LIZARD L1_AC11 | FROG L1-35 | ZEBRAFISH L1-12A |
| HYRAX | LIZARD L1_AC12 | FROG L1-38 | ZEBRAFISH L1-12B |
| LEMUR | LIZARD L1_AC14 | FROG L1-39 | ZEBRAFISH L1-13A |
| OPOSSUM | LIZARD L1_AC15 | FROG L1-46 | ZEBRAFISH L1-13B |
| PANDA | LIZARD L1_AC17 | FROG L1-47 | ZEBRAFISH L1-13C |
| RABBIT | LIZARD L1_AC18 | ZEBRAFISH L1-1 | ZEBRAFISH L1-13D |
| RAT | LIZARD L1_AC20 | ZEBRAFISH L1-1B | ZEBRAFISH L1-16B |
| HORSE | FROG L1-6 | ZEBRAFISH L1-1D | ZEBRAFISH L1-17B |
| MOUSE | FROG L1-11 | ZEBRAFISH L1-6 | |

#### Supplemental Figure 5

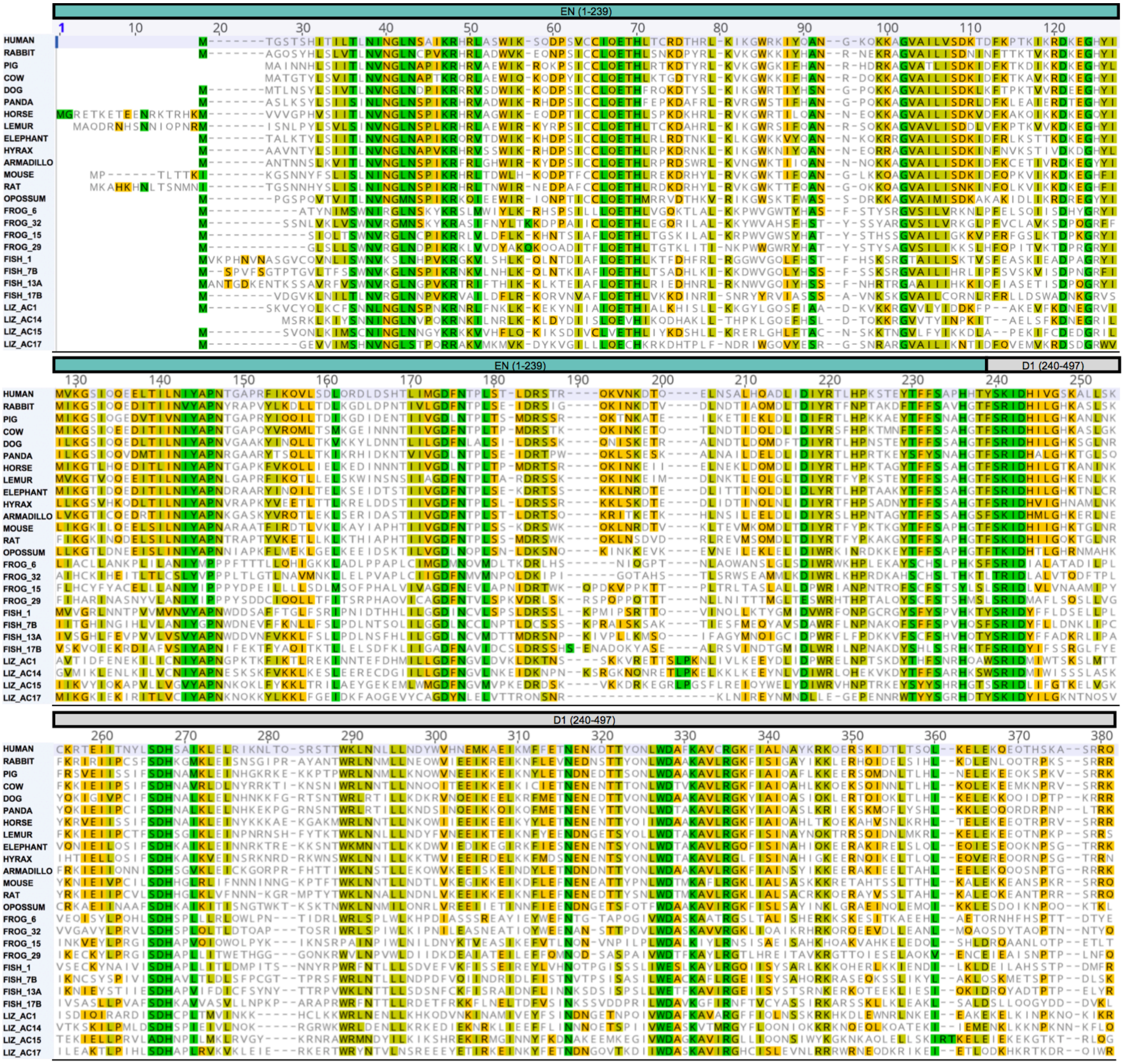

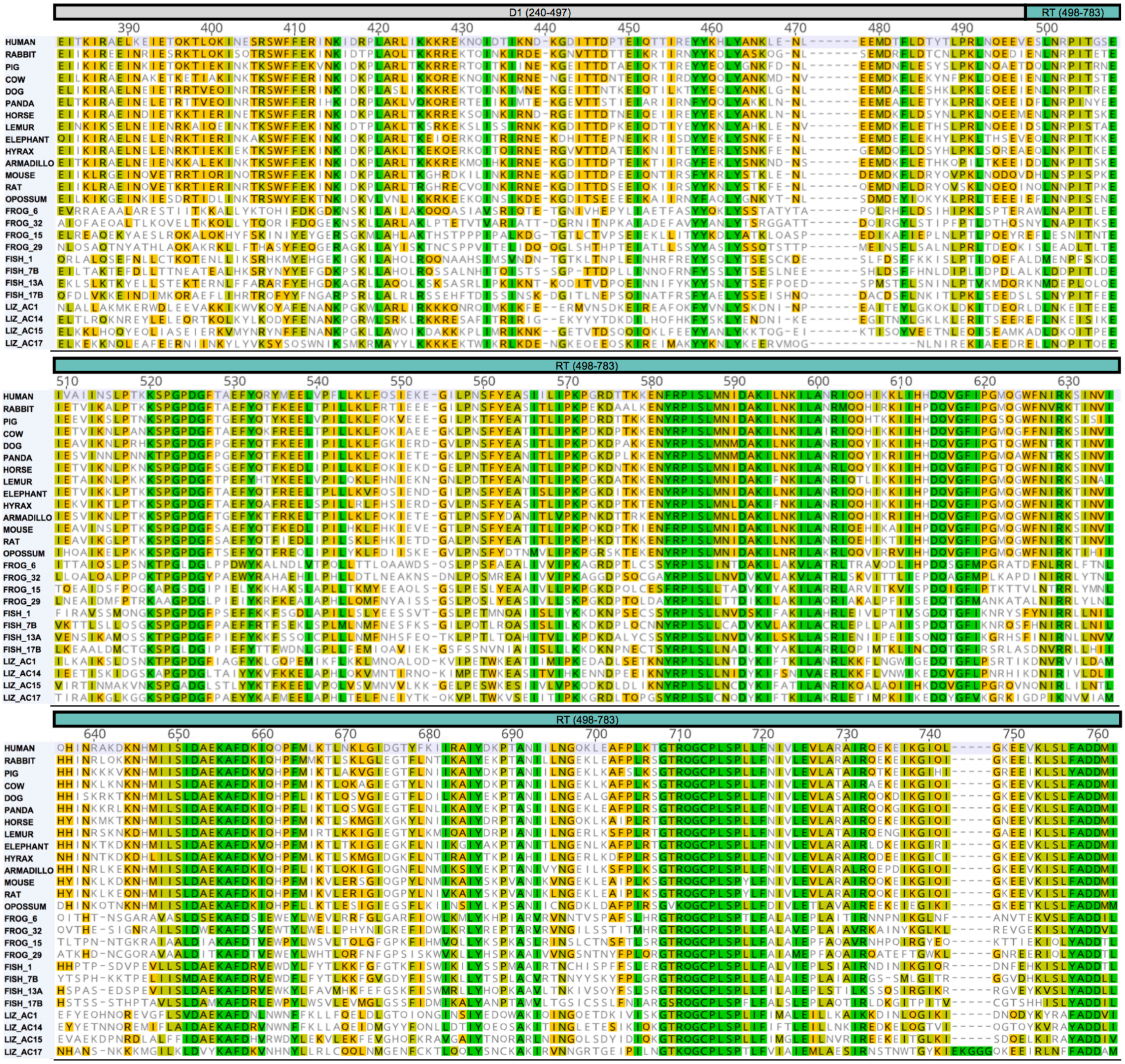
D1 (240-497)
RT (498-783)
HUMAN
RABBIT
PIG
COW
DOG
PANDA
HORSE
LEMUR
ELEPHANT
HYRAX
ARMADILLO
MOUSE
RAT
OPOSSUM
FROG_6
FROG_32
FROG_15
FROG_29
FISH_1
FISH_7B
FISH_13A
FISH_17B
LIZ_AC1
LIZ_AC14
LIZ_AC15
LIZ_AC17

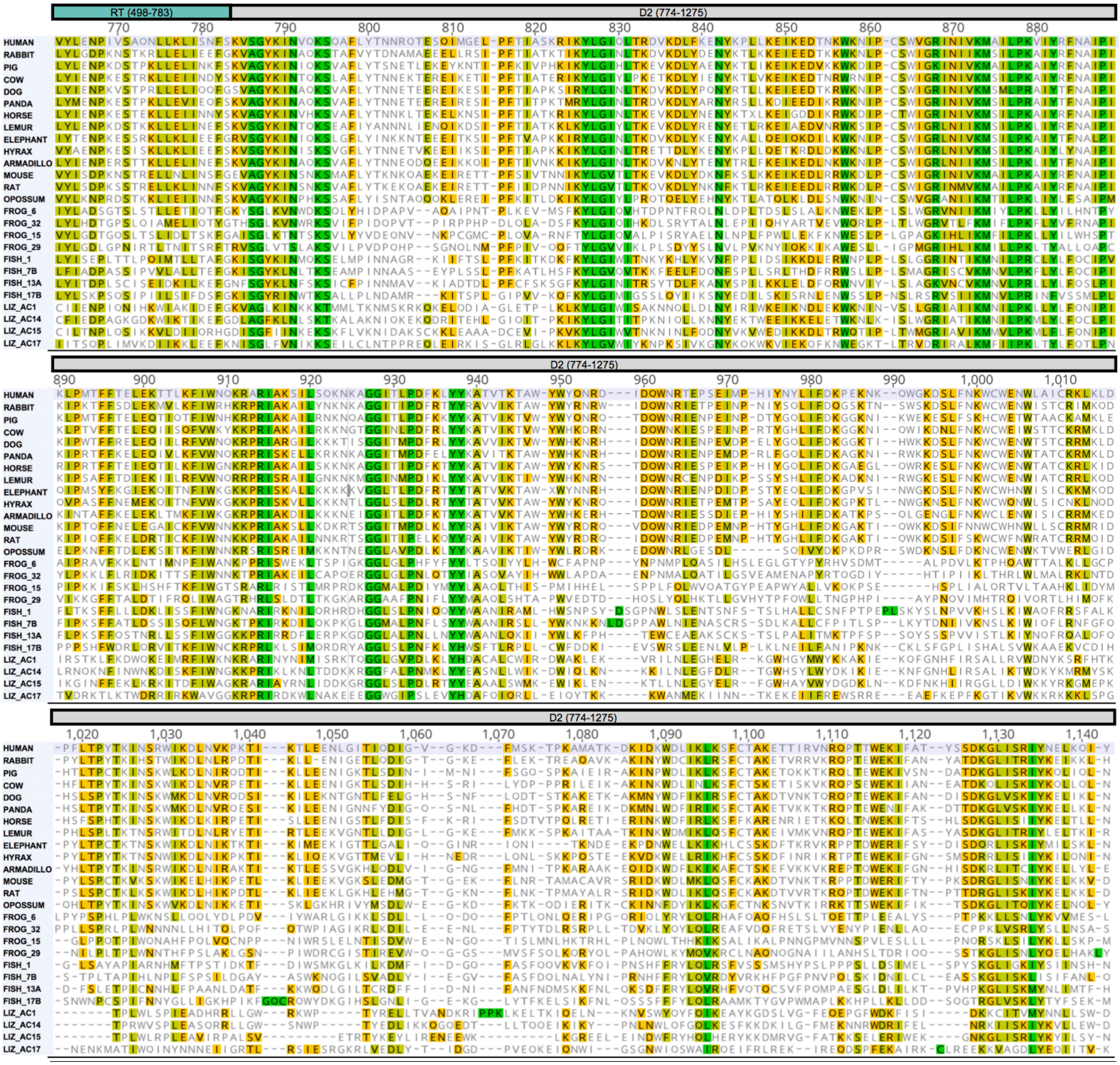
RT (498-783)
D2 (774-1275)
770 780 790 800 810 820 830 840 850 860 870 880
890 900 910 920 930 940 950 960 970 980 990 1,000 1,010
1,020 1,030 1,040 1,050 1,060 1,070 1,080 1,090 1,100 1,110 1,120 1,130 1,140
HUMAN
RABBIT
PIG
COW
DOG
PANDA
HORSE
LEMUR
ELEPHANT
HYRAX
ARMADILLO
MOUSE
RAT
OPOSSUM
FROG_6
FROG_32
FROG_15
FROG_29
FISH_1
FISH_7B
FISH_13A
FISH_17B
LIZ_AC1
LIZ_AC14
LIZ_AC15
LIZ_AC17

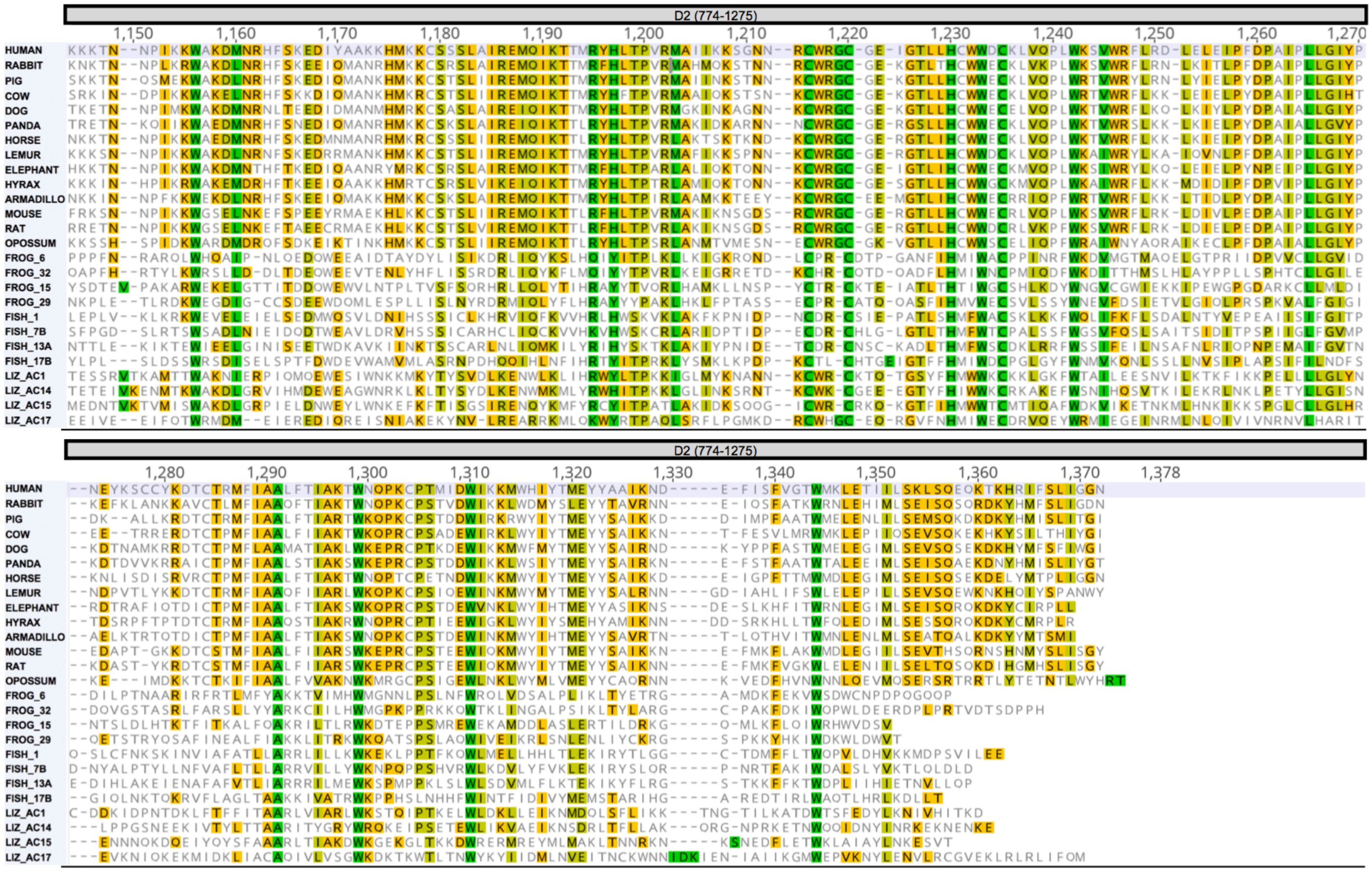
D2 (774-1275)
1,150
1,160
1,170
1,180
1,190
1,200
1,210
1,220
1,230
1,240
1,250
1,260
1,270
HUMAN
RABBIT
PIG
COW
DOG
PANDA
HORSE
LEMUR
ELEPHANT
HYRAX
ARMADILLO
MOUSE
RAT
OPOSSUM
FROG_6
FROG_32
FROG_15
FROG_29
FISH_1
FISH_7B
FISH_13A
FISH_17B
LIZ_AC1
LIZ_AC14
LIZ_AC15
LIZ_AC17
D2 (774-1275)
1,280
1,290
1,300
1,310
1,320
1,330
1,340
1,350
1,360
1,370
1,378

**Supplemental Table 8**

| Mutant ID | Conservation (residue with highest identity value as representative for 3xala mutant) | | | Avg retroT | Domain |
|---|---|---|---|---|---|
| | Mammals | Mammals + Verts | | | |
| pEA1362_2_3_TG | 14 | 4 | | 25 | EN |
| pEA920_4_6_STS | 20 | 6 | | 72 | EN |
| pEA921_7_9_HIT | 62 | 29 | | 18 | EN |
| pEA922_10_12_ILT | 56 | 52 | | 4 | EN |
| pEA923_13_15_LNI | 100 | 100 | | 6 | EN |
| pEA924_16_18_NGL | 100 | 96 | | 5 | EN |
| pEA925_19_21_NSA | 100 | 67 | | 9 | EN |
| pEA926_22_24_IKR | 100 | 96 | | 11 | EN |
| pEA927_25_27_HRL | 86 | 27 | | 36 | EN |
| pEA928_28_30_ASW | 100 | 14 | | 18 | EN |
| pEA929_31_33_IKS | 73 | 42 | | 10 | EN |
| pEA930_34_36_QDP | 100 | 20 | | 43 | EN |
| pEA931_37_39_SVC | 100 | 66 | | 0 | EN |
| pEA932_40_42_CIQ | 100 | 93 | | 2 | EN |
| pEA933_43_45_ETH | 100 | 100 | | 2 | EN |
| pEA934_46_48_LTC | 62 | 45 | | 6 | EN |
| pEA935_49_51_RDT | 100 | 31 | | 81 | EN |
| pEA936_52_54_HRL | 86 | 68 | | 4 | EN |
| pEA937_55_57_KIK | 73 | 27 | | 15 | EN |
| pEA938_58_60_GWR | 100 | 24 | | 4 | EN |
| pEA939_61_63_KIY | 62 | 37 | | 0 | EN |
| pEA940_64_66_QAN | 86 | 46 | | 84 | EN |
| pEA941_67_69_GKQ | 41 | 13 | | 52 | EN |
| pEA942_70_72_KKA | 86 | 50 | | 7 | EN |
| pEA943_73_75_GVA | 100 | 100 | | 4 | EN |
| pEA944_76_78_ILV | 86 | 60 | | 2 | EN |
| pEA945_79_81_SDK | 100 | 28 | | 2 | EN |
| pEA946_82_84_TDF | 86 | 51 | | 6 | EN |
| pEA947_85_87_KPT | 42 | 14 | | 18 | EN |
| pEA948_88_90_KIK | 39 | 18 | | 10 | EN |
| pEA949_91_93_RDK | 100 | 100 | | 10 | EN |
| pEA950_94_96_EGH | 100 | 100 | | 3 | EN |
| pEA951_97_99_YIM | 74 | 36 | | 2 | EN |
| pEA952_100_102_VKG | 100 | 42 | | 3 | EN |
| pEA953_103_105_SIQ | 52 | 37 | | 28 | EN |
| pEA954_106_108_QEE | 60 | 16 | | 33 | EN |
| pEA955_109_111_LTI | 74 | 38 | | 5 | EN |
| pEA956_112_114_LNI | 100 | 70 | | 2 | EN |
| pEA957_115_117_YAP | 100 | 100 | | 2 | EN |
| pEA958_118_120_NTG | 100 | 70 | | 24 | EN |
| pEA959_121_123_APR | 100 | 15 | | 65 | EN |
| pEA960_124_126_FIK | 51 | 37 | | 5 | EN |
| pEA961_127_129_QVL | 100 | 26 | | 2 | EN |
| pEA962_130_132_SDL | 51 | 38 | | 40 | EN |
| pEA963_133_135_QRD | 62 | 12 | | 23 | EN |
| pEA964_136_138_LDS | 56 | 9 | | 7 | EN |
| pEA965_139_141_HTL | 86 | 64 | | 3 | EN |
| pEA966_142_144_IMG | 100 | 96 | | 2 | EN |
| pEA967_145_147_DFN | 100 | 100 | | 4 | EN |
| pEA968_148_150_TPL | 100 | 39 | | 6 | EN |
| pEA969_151_153_STL | 56 | 24 | | 64 | EN |
| pEA970_154_156_DRS | 100 | 93 | | 2 | EN |
| pEA971_157_159_TRQ | 56 | 17 | | 52 | EN |
| pEA972_160_162_KVN | 60 | 11 | | 48 | EN |
| pEA973_163_165_KDT | 62 | 22 | | 97 | EN |
| pEA974_166_168_QEL | 86 | 31 | | 19 | EN |
| pEA975_169_171_NSA | 41 | 13 | | 79 | EN |
| pEA976_172_174_LHQ | 52 | 17 | | 25 | EN |
| pEA977_175_177_ADL | 86 | 54 | | 8 | EN |
| pEA978_178_180_IDI | 100 | 96 | | 3 | EN |

| Mutant ID | Conservation (residue with highest identity value as representative for 3xala mutant) | | Avg retroT | Domain |
|---|---|---|---|---|
| | Mammals | Mammals + Verts | | |
| pEA979_181_183_YRT | 100 | 86 | 3 | EN |
| pEA980_184_186_LHP | 86 | 73 | 5 | EN |
| pEA981_187_189_KST | 25 | 33 | 88 | EN |
| pEA982_190_192_EYT | 86 | 54 | 4 | EN |
| pEA983_193_195_FFS | 100 | 93 | 2 | EN |
| pEA984_196_198_APH | 100 | 76 | 10 | EN |
| pEA985_199_201_HTY | 86 | 33 | 6 | EN |
| pEA986_202_204_SKI | 100 | 96 | 7 | EN |
| pEA987_205_207_DHI | 100 | 100 | 2 | EN |
| pEA988_208_210_VGS | 100 | 36 | 7 | EN |
| pEA989_211_213_KAL | 73 | 23 | 6 | EN |
| pEA990_214_216_LSK | 73 | 21 | 61 | EN |
| pEA991_217_219_CKR | 28 | 19 | 92 | EN |
| pEA992_220_222_TEI | 86 | 42 | 46 | EN |
| pEA993_223_225_ITN | 52 | 31 | 33 | EN |
| pEA994_226_228_YLS | 100 | 86 | 17 | EN |
| pEA995_229_231_DHS | 100 | 100 | 3 | EN |
| pEA996_232_234_AIK | 74 | 43 | 14 | EN |
| pEA997_235_237_LEL | 73 | 36 | 7 | EN |
| pEA998_238_240_RIK | 51 | 17 | 91 | EN (1-239) |
| pEA999_241_243_NLT | 15 | 8 | 104 | DESERT 1 (240-379) |
| pEA1000_244_246_QSR | 21 | 11 | 60 | DESERT 1 |
| pEA1001_247_249_STT | 32 | 11 | 88 | DESERT 1 |
| pEA1002_250_252_WKL | 100 | 96 | 5 | DESERT 1 |
| pEA1003_253_255_NNL | 74 | 49 | 42 | DESERT 1 |
| pEA1004_256_258_LLN | 100 | 67 | 10 | DESERT 1 |
| pEA1005_259_261_DYW | 26 | 21 | 29 | DESERT 1 |
| pEA1006_262_264_VHN | 64 | 20 | 106 | DESERT 1 |
| pEA1007_265_267_EMK | 86 | 32 | 22 | DESERT 1 |
| pEA1008_268_270_AEI | 74 | 53 | 13 | DESERT 1 |
| pEA1009_271_273_KMF | 62 | 47 | 7 | DESERT 1 |
| pEA1010_274_276_FET | 86 | 23 | 16 | DESERT 1 |
| pEA1011_277_279_NEN | 100 | 96 | 12 | DESERT 1 |
| pEA1012_280_282_KDT | 100 | 28 | 28 | DESERT 1 |
| pEA1013_283_285_TYQ | 73 | 28 | 19 | DESERT 1 |
| pEA1014_286_288_NLW | 100 | 100 | 3 | DESERT 1 |
| pEA1015_289_291_DAF | 100 | 51 | 3 | DESERT 1 |
| pEA1016_292_294_KAV | 100 | 100 | 3 | DESERT 1 |
| pEA1017_295_297_CRG | 100 | 100 | 5 | DESERT 1 |
| pEA1018_298_300_KFI | 100 | 73 | 3 | DESERT 1 |
| pEA1019_301_303_ALN | 56 | 29 | 103 | DESERT 1 |
| pEA1020_304_306_AYK | 86 | 29 | 73 | DESERT 1 |
| pEA1021_307_309_RKQ | 86 | 58 | 62 | DESERT 1 |
| pEA1022_310_312_ERS | 73 | 14 | 64 | DESERT 1 |
| pEA1023_313_315_KID | 42 | 18 | 62 | DESERT 1 |
| pEA1024_316_318_TLT | 100 | 57 | 59 | DESERT 1 |
| pEA1025_319_321_SQL | 74 | 36 | 73 | DESERT 1 |
| pEA1026_322_324_KEL | 73 | 38 | 69 | DESERT 1 |
| pEA1027_325_327_EKQ | 100 | 52 | 87 | DESERT 1 |
| pEA1028_328_330_EQT | 46 | 17 | 77 | DESERT 1 |
| pEA1029_331_333_HSK | 86 | 56 | 83 | DESERT 1 |
| pEA1030_334_336_ASR | 74 | 15 | 94 | DESERT 1 |
| pEA1031_337_339_RQE | 86 | 32 | 20 | DESERT 1 |
| pEA1032_340_342_ITK | 100 | 31 | 31 | DESERT 1 |
| pEA1033_343_345_IRA | 74 | 24 | 30 | DESERT 1 |
| pEA1034_346_348_ELK | 100 | 31 | 14 | DESERT 1 |
| pEA1035_349_351_EIE | 100 | 23 | 59 | DESERT 1 |
| pEA1036_352_354_TQK | 39 | 17 | 54 | DESERT 1 |
| pEA1037_355_357_TLQ | 74 | 25 | 31 | DESERT 1 |
| pEA1038_358_360_KIN | 100 | 28 | 57 | DESERT 1 |

| Mutant ID | Conservation (residue with highest identity value as representative for 3xala mutant) | | Avg retroT | Domain |
|---|---|---|---|---|
| | Mammals | Mammals + Verts | | |
| pEA1039_361_363_ESR | 73 | 40 | 31 | DESERT 1 |
| pEA1040_364_366_SWF | 100 | 25 | 12 | DESERT 1 |
| pEA1041_367_369_FER | 100 | 56 | 24 | DESERT 1 |
| pEA1042_370_372_INK | 100 | 86 | 17 | DESERT 1 |
| pEA1043_373_375_IDR | 100 | 48 | 61 | DESERT 1 |
| pEA1044_376_378_PLA | 100 | 90 | 42 | DESERT 1 |
| pEA1045_379_381_RLI | 100 | 27 | 65 | DESERT 1 (240-379) |
| pEA1046_382_384_KKK | 73 | 46 | 57 | Z (380-480) |
| pEA1047_385_387_REK | 86 | 22 | 59 | Z |
| pEA1048_388_390_NQI | 100 | 80 | 60 | Z |
| pEA1049_391_393_DTI | 100 | 74 | 60 | Z |
| pEA1050_394_396_KND | 62 | 22 | 77 | Z |
| pEA1051_397_399_KGD | 86 | 56 | 55 | Z |
| pEA1052_400_402_ITT | 100 | 26 | 22 | Z |
| pEA1053_403_405_DPT | 51 | 27 | 45 | Z |
| pEA1054_406_408_EIQ | 100 | 96 | 14 | Z |
| pEA1055_409_411_TTI | 86 | 50 | 28 | Z |
| pEA1056_412_414_REY | 74 | 50 | 28 | Z |
| pEA1057_415_417_YKH | 74 | 86 | 33 | Z |
| pEA1058_418_420_LYA | 100 | 100 | 31 | Z |
| pEA1059_421_423_NKL | 100 | 21 | 66 | Z |
| pEA1060_424_426_ENL | 100 | 25 | 50 | Z |
| pEA1061_427_429_EEM | 100 | 24 | 38 | Z |
| pEA1062_430_432_DTF | 86 | 51 | 20 | Z |
| pEA1063_433_435_LDT | 86 | 55 | 6 | Z |
| pEA1064_436_438_YTL | 51 | 47 | 18 | Z |
| pEA1065_439_441_PRL | 100 | 54 | 23 | Z |
| pEA1066_442_444_NQE | 60 | 26 | 73 | Z |
| pEA1067_445_447_EVE | 62 | 23 | 43 | Z |
| pEA1068_448_450_SLN | 100 | 80 | 30 | Z |
| pEA1069_451_453_RPI | 100 | 74 | 31 | Z |
| pEA1070_454_456_TGS | 53 | 43 | 88 | Z |
| pEA1071_457_459_EIV | 100 | 96 | 3 | Z |
| pEA1072_460_462_AII | 100 | 76 | 16 | Z |
| pEA1073_463_465_NSL | 100 | 53 | 13 | Z |
| pEA1074_466_468_PTK | 100 | 28 | 40 | Z |
| pEA1075_469_471_KSP | 100 | 100 | 62 | Z |
| pEA1076_472_474_GPD | 100 | 100 | 4 | Z |
| pEA1077_475_477_GFT | 100 | 100 | 7 | Z |
| pEA1078_478_480_AEF | 100 | 54 | 18 | Z (380-480) |
| pEA1079_481_483_YQR | 100 | 90 | 14 | |
| pEA1080_484_486_YKE | 86 | 44 | 8 | |
| pEA1081_487_489_ELV | 74 | 56 | 10 | |
| pEA1082_490_492_PFL | 100 | 83 | 8 | |
| pEA1083_493_495_LKL | 86 | 26 | 6 | |
| pEA1084_496_498_FQS | 100 | 44 | 28 | RT (498-773) |
| pEA1085_499_501_IEK | 86 | 24 | 79 | RT |
| pEA1086_502_504_EGI | 100 | 35 | 38 | RT |
| pEA1087_505_507_LPN | 100 | 83 | 8 | RT |
| pEA1088_508_510_SFY | 100 | 41 | 9 | RT |
| pEA1089_511_513_EAS | 86 | 74 | 41 | RT |
| pEA1090_514_516_IIL | 100 | 96 | 6 | RT |
| pEA1091_517_519_IPK | 100 | 100 | 8 | RT |
| pEA1092_520_522_PGR | 100 | 54 | 10 | RT |
| pEA1093_523_525_DTT | 86 | 76 | 80 | RT |
| pEA1094_526_528_KKE | 100 | 27 | 83 | RT |
| pEA1095_529_531_NFR | 100 | 100 | 6 | RT |
| pEA1096_532_534_PIS | 100 | 100 | 6 | RT |
| pEA1097_535_537_LMN | 100 | 89 | 10 | RT |
| pEA1098_538_540_IDA | 100 | 96 | 8 | RT |

233

| Mutant ID | Conservation (residue with highest identity value as representative for 3xala mutant) | | Avg retroT | Domain |
|---|---|---|---|---|
| | Mammals | Mammals + Verts | | |
| pEA1099_541_543_KIL | 100 | 96 | 4 | RT |
| pEA1100_544_546_NKI | 100 | 67 | 58 | RT |
| pEA1101_547_549_LAN | 100 | 89 | 53 | RT |
| pEA1102_550_552_RIQ | 100 | 100 | 8 | RT |
| pEA1103_553_555_QHI | 86 | 38 | 89 | RT |
| pEA1104_556_558_KKL | 74 | 34 | 43 | RT |
| pEA1105_559_561_IHH | 100 | 67 | 16 | RT |
| pEA1106_562_564_DQV | 100 | 100 | 6 | RT |
| pEA1107_565_567_GFI | 100 | 100 | 20 | RT |
| pEA1108_568_570_PGM | 100 | 49 | 53 | RT |
| pEA1109_571_573_QGW | 100 | 22 | 16 | RT |
| pEA1110_574_576_FNI | 100 | 90 | 36 | RT |
| pEA1111_577_579_RKS | 100 | 93 | 5 | RT |
| pEA1112_580_582_INV | 100 | 30 | 30 | RT |
| pEA1113_583_585_IQH | 100 | 42 | 61 | RT |
| pEA1114_586_588_INR | 100 | 14 | 57 | RT |
| pEA1115_589_591_AKD | 60 | 17 | 79 | RT |
| pEA1116_592_594_KNH | 100 | 16 | 54 | RT |
| pEA1117_595_597_MII | 100 | 32 | 6 | RT |
| pEA1118_598_600_SID | 100 | 100 | 5 | RT |
| pEA1119_601_603_AEK | 100 | 100 | 62 | RT |
| pEA1120_604_606_AFD | 100 | 100 | 4 | RT |
| pEA1121_607_609_KIQ | 100 | 40 | 39 | RT |
| pEA1122_610_612_QPF | 100 | 53 | 5 | RT |
| pEA1123_613_615_MLK | 86 | 46 | 6 | RT |
| pEA1124_616_618_TLN | 100 | 56 | 35 | RT |
| pEA1125_619_621_KLG | 100 | 46 | 13 | RT |
| pEA1126_622_624_IDG | 100 | 38 | 8 | RT |
| pEA1127_625_627_TYF | 86 | 62 | 19 | RT |
| pEA1128_628_630_KII | 73 | 46 | 4 | RT |
| pEA1129_631_633_RAI | 100 | 46 | 25 | RT |
| pEA1130_634_636_YDK | 100 | 96 | 14 | RT |
| pEA1131_637_639_PTA | 100 | 96 | 63 | RT |
| pEA1132_640_642_NII | 100 | 48 | 4 | RT |
| pEA1133_643_645_LNG | 100 | 86 | 13 | RT |
| pEA1134_646_648_QKL | 100 | 37 | 45 | RT |
| pEA1135_649_651_EAF | 64 | 55 | 6 | RT |
| pEA1136_652_654_PLK | 86 | 44 | 35 | RT |
| pEA1137_655_657_TGT | 100 | 96 | 11 | RT |
| pEA1138_658_660_RQG | 100 | 100 | 9 | RT |
| pEA1139_661_663_CPL | 100 | 100 | 6 | RT |
| pEA1140_664_666_SPL | 100 | 100 | 5 | RT |
| pEA1141_667_669_LFN | 100 | 93 | 12 | RT |
| pEA1142_670_672_IVL | 100 | 39 | 14 | RT |
| pEA1143_673_675_EVL | 100 | 100 | 5 | RT |
| pEA1144_676_678_ARA | 100 | 53 | 123 | RT |
| pEA1145_679_681_IRQ | 100 | 77 | 6 | RT |
| pEA1146_682_684_EKE | 60 | 16 | 29 | RT |
| pEA1147_685_687_IKG | 100 | 93 | 4 | RT |
| pEA1148_688_690_IQL | 100 | 39 | 9 | RT |
| pEA1149_691_693_GKE | 100 | 25 | 52 | RT |
| pEA1150_694_696_EVK | 100 | 74 | 3 | RT |
| pEA1151_697_699_LSL | 100 | 58 | 37 | RT |
| pEA1152_700_702_FAD | 100 | 100 | 9 | RT |
| pEA1153_703_705_DMI | 100 | 100 | 4 | RT |
| pEA1154_706_708_VYL | 100 | 39 | 7 | RT |
| pEA1155_709_711_ENP | 100 | 80 | 16 | RT |
| pEA1156_712_714_IVS | 100 | 46 | 34 | RT |
| pEA1157_715_717_AQN | 60 | 25 | 88 | RT |
| pEA1158_718_720_LLK | 100 | 34 | 7 | RT |

234

| Mutant ID | Conservation (residue with highest identity value as representative for 3xala mutant) | | | Avg retroT | Domain |
|---|---|---|---|---|---|
| | Mammals | Mammals + Verts | | | |
| pEA1159_721_723_LIS | 100 | 51 | | 6 | RT |
| pEA1160_724_726_NFS | 74 | 56 | | 5 | RT |
| pEA1161_727_729_KVS | 100 | 57 | | 20 | RT |
| pEA1162_730_732_GYK | 100 | 93 | | 6 | RT |
| pEA1163_733_735_INV | 100 | 96 | | 11 | RT |
| pEA1164_736_738_QKS | 100 | 100 | | 5 | RT |
| pEA1165_739_741_QAF | 100 | 33 | | 8 | RT |
| pEA1166_742_744_LYT | 100 | 33 | | 6 | RT |
| pEA1167_745_747_NNR | 73 | 12 | | 60 | RT |
| pEA1168_748_750_QTE | 62 | 12 | | 92 | RT |
| pEA1169_751_753_SQI | 62 | 21 | | 120 | RT |
| pEA1170_754_756_MGE | 31 | 9 | | 116 | RT |
| pEA1171_757_759_LPF | 100 | 34 | | 22 | RT |
| pEA1172_760_762_TIA | 86 | 23 | | 92 | RT |
| pEA1173_763_765_SKR | 41 | 22 | | 66 | RT |
| pEA1174_766_768_IKY | 100 | 100 | | 10 | RT |
| pEA1175_769_771_LGI | 100 | 100 | | 4 | RT |
| pEA1176_772_774_QLT | 100 | 36 | | 20 | RT (498-773) |
| pEA1177_775_777_RDV | 64 | 16 | | 78 | DESERT 2 (774-1275) |
| pEA1178_778_780_KDL | 100 | 52 | | 44 | DESERT 2 |
| pEA1179_781_783_FKE | 86 | 19 | | 83 | DESERT 2 |
| pEA1180_784_786_NYK | 100 | 100 | | 14 | DESERT 2 |
| pEA1181_787_789_PLL | 86 | 30 | | 42 | DESERT 2 |
| pEA1182_790_792_KEI | 73 | 37 | | 54 | DESERT 2 |
| pEA1183_793_795_KEE | 100 | 35 | | 75 | DESERT 2 |
| pEA1184_796_798_TNK | 40 | 32 | | 104 | DESERT 2 |
| pEA1185_799_801_WKN | 100 | 100 | | 62 | DESERT 2 |
| pEA1186_802_804_IPC | 100 | 47 | | 80 | DESERT 2 |
| pEA1187_805_807_SWV | 100 | 65 | | 25 | DESERT 2 |
| pEA1188_808_810_GRI | 100 | 77 | | 34 | DESERT 2 |
| pEA1189_811_813_NIV | 100 | 35 | | 41 | DESERT 2 |
| pEA1190_814_816_KMA | 100 | 100 | | 35 | DESERT 2 |
| pEA1191_817_819_ILP | 100 | 96 | | 11 | DESERT 2 |
| pEA1192_820_822_KVI | 100 | 64 | | 19 | DESERT 2 |
| pEA1193_823_825_YRF | 100 | 80 | | 100 | DESERT 2 |
| pEA1194_826_828_NAI | 100 | 35 | | 56 | DESERT 2 |
| pEA1195_829_831_PIK | 100 | 96 | | 5 | DESERT 2 |
| pEA1196_832_834_LPM | 86 | 42 | | 66 | DESERT 2 |
| pEA1197_835_837_TFF | 100 | 73 | | 35 | DESERT 2 |
| pEA1198_838_840_TEL | 53 | 29 | | 67 | DESERT 2 |
| pEA1199_841_843_EKT | 86 | 23 | | 82 | DESERT 2 |
| pEA1200_844_846_TLK | 52 | 31 | | 98 | DESERT 2 |
| pEA1201_847_849_FIW | 100 | 89 | | 8 | DESERT 2 |
| pEA1202_850_852_NQK | 86 | 70 | | 83 | DESERT 2 |
| pEA1203_853_855_RAR | 100 | 86 | | 19 | DESERT 2 |
| pEA1204_856_858_IAK | 100 | 60 | | 52 | DESERT 2 |
| pEA1205_859_861_SIL | 86 | 68 | | 56 | DESERT 2 |
| pEA1206_862_864_SQK | 86 | 19 | | 73 | DESERT 2 |
| pEA1207_865_867_NKA | 42 | 24 | | 122 | DESERT 2 |
| pEA1208_868_870_GGI | 100 | 100 | | 60 | DESERT 2 |
| pEA1209_871_873_TLP | 100 | 100 | | 69 | DESERT 2 |
| pEA1210_874_876_DFK | 86 | 38 | | 7 | DESERT 2 |
| pEA1211_877_879_LYY | 100 | 100 | | 6 | DESERT 2 |
| pEA1212_880_882_KAT | 100 | 83 | | 60 | DESERT 2 |
| pEA1213_883_885_VTK | 100 | 34 | | 62 | DESERT 2 |
| pEA1214_886_888_TAW | 100 | 20 | | 36 | DESERT 2 |
| pEA1215_889_891_YWY | 100 | 70 | | 4 | DESERT 2 |
| pEA1216_892_894_QNR | 100 | 13 | | 38 | DESERT 2 |
| pEA1217_895_897_DID | 100 | 12 | | 55 | DESERT 2 |
| pEA1218_898_900_QWN | 100 | 49 | | 60 | DESERT 2 |

235

| Mutant ID | Conservation (residue with highest identity value as representative for 3xala mutant) | | | Avg retroT | Domain |
|---|---|---|---|---|---|
| | Mammals | Mammals + Verts | | | |
| pEA1219_901_903_RTE | 86 | 83 | | 69 | DESERT 2 |
| pEA1220_904_906_PSE | 64 | 13 | | 120 | DESERT 2 |
| pEA1221_907_909_IMP | 86 | 19 | | 112 | DESERT 2 |
| pEA1222_910_912_HIY | 73 | 19 | | 88 | DESERT 2 |
| pEA1223_913_915_NYL | 74 | 26 | | 105 | DESERT 2 |
| pEA1224_916_918_IFD | 100 | 34 | | 7 | DESERT 2 |
| pEA1225_919_921_KPE | 86 | 19 | | 95 | DESERT 2 |
| pEA1226_922_924_KNK | 60 | 20 | | 79 | DESERT 2 |
| pEA1227_925_927_QWG | 86 | 9 | | 112 | DESERT 2 |
| pEA1228_928_930_KDS | 86 | 18 | | 73 | DESERT 2 |
| pEA1229_931_933_LFN | 100 | 22 | | 49 | DESERT 2 |
| pEA1230_934_936_KWC | 100 | 27 | | 75 | DESERT 2 |
| pEA1231_937_939_WEN | 86 | 19 | | 64 | DESERT 2 |
| pEA1232_940_942_WLA | 100 | 79 | | 19 | DESERT 2 |
| pEA1233_943_945_ICR | 86 | 16 | | 96 | DESERT 2 |
| pEA1234_946_948_KLK | 51 | 21 | | 97 | DESERT 2 |
| pEA1235_949_951_LDP | 86 | 14 | | 71 | DESERT 2 |
| pEA1236_952_954_FLT | 86 | 23 | | 16 | DESERT 2 |
| pEA1237_955_957_PYT | 100 | 40 | | 4 | DESERT 2 |
| pEA1238_958_960_KIN | 100 | 46 | | 8 | DESERT 2 |
| pEA1239_961_963_SRW | 100 | 30 | | 10 | DESERT 2 |
| pEA1240_964_966_IKD | 86 | 22 | | 42 | DESERT 2 |
| pEA1241_967_969_LNV | 100 | 16 | | 8 | DESERT 2 |
| pEA1242_970_972_KPK | 47 | 13 | | 64 | DESERT 2 |
| pEA1243_973_975_TIK | 74 | 16 | | 14 | DESERT 2 |
| pEA1244_976_978_TLE | 60 | 47 | | 28 | DESERT 2 |
| pEA1245_979_981_ENL | 62 | 22 | | 63 | DESERT 2 |
| pEA1246_982_984_GIT | 86 | 22 | | 25 | DESERT 2 |
| pEA1247_985_987_IQD | 51 | 34 | | 51 | DESERT 2 |
| pEA1248_988_990_IGV | 35 | 29 | | 10 | DESERT 2 |
| pEA1249_991_993_GKD | 34 | 14 | | 97 | DESERT 2 |
| pEA1250_994_996_FMS | 53 | 30 | | 8 | DESERT 2 |
| pEA1251_997_999_KTP | 31 | 27 | | 93 | DESERT 2 |
| pEA1252_1000_1002_KAM | 40 | 21 | | 103 | DESERT 2 |
| pEA1253_1003_1005_ATK | 28 | 15 | | 109 | DESERT 2 |
| pEA1254_1006_1008_DKI | 53 | 20 | | 77 | DESERT 2 |
| pEA1255_1009_1011_DKW | 86 | 23 | | 11 | DESERT 2 |
| pEA1256_1012_1014_DLI | 74 | 45 | | 11 | DESERT 2 |
| pEA1257_1015_1017_KLK | 53 | 48 | | 11 | DESERT 2 |
| pEA1258_1018_1020_SFC | 86 | 21 | | 6 | DESERT 2 |
| pEA1259_1021_1023_TAK | 86 | 17 | | 93 | DESERT 2 |
| pEA1260_1024_1026_ETT | 42 | 9 | | 91 | DESERT 2 |
| pEA1261_1027_1029_IRV | 42 | 8 | | 110 | DESERT 2 |
| pEA1262_1030_1032_NRQ | 86 | 14 | | 72 | DESERT 2 |
| pEA1263_1033_1035_PTT | 86 | 24 | | 126 | DESERT 2 |
| pEA1264_1036_1038_WEK | 100 | 42 | | 29 | DESERT 2 |
| pEA1265_1039_1041_IFA | 100 | 34 | | 8 | DESERT 2 |
| pEA1266_1042_1044_TYS | 26 | 10 | | 106 | DESERT 2 |
| pEA1267_1045_1047_SDK | 100 | 42 | | 41 | DESERT 2 |
| pEA1268_1048_1050_GLI | 100 | 44 | | 15 | DESERT 2 |
| pEA1269_1051_1053_SRI | 100 | 60 | | 55 | DESERT 2 |
| pEA1270_1054_1056_YNE | 100 | 96 | | 66 | DESERT 2 |
| pEA1271_1057_1059_LKQ | 86 | 44 | | 82 | DESERT 2 |
| pEA1272_1060_1062_IYK | 53 | 12 | | 67 | DESERT 2 |
| pEA1273_1063_1065_KKT | 53 | 12 | | 108 | DESERT 2 |
| pEA1274_1066_1068_NNP | 86 | 14 | | 57 | DESERT 2 |
| pEA1275_1069_1071_IKK | 74 | 29 | | 43 | DESERT 2 |
| pEA1276_1072_1074_WAK | 100 | 100 | | 18 | DESERT 2 |
| pEA1277_1075_1077_DMN | 74 | 35 | | 5 | DESERT 2 |
| pEA1278_1078_1080_RHF | 86 | 27 | | 14 | DESERT 2 |
| pEA1279_1081_1083_SKE | 74 | 35 | | 94 | DESERT 2 |
| pEA1280_1084_1086_DIY | 50 | 51 | | 65 | DESERT 2 |

236

| Mutant ID | Conservation (residue with highest identity value as representative for 3xala mutant) | | Avg retroT | Domain |
|---|---|---|---|---|
| | Mammals | Mammals + Verts | | |
| pEA1281_1087_1089_AAK | 86 | 21 | 64 | DESERT 2 |
| pEA1282_1090_1092_KHM | 86 | 14 | 15 | DESERT 2 |
| pEA1283_1093_1095_KKC | 100 | 24 | 65 | DESERT 2 |
| pEA1284_1096_1098_SSS | 100 | 33 | 44 | DESERT 2 |
| pEA1285_1099_1101_LAI | 100 | 20 | 50 | DESERT 2 |
| pEA1286_1102_1104_REM | 100 | 41 | 24 | DESERT 2 |
| pEA1287_1105_1107_QIK | 100 | 89 | 28 | DESERT 2 |
| pEA1288_1108_1110_TTM | 100 | 24 | 44 | DESERT 2 |
| pEA1289_1111_1113_RYH | 100 | 70 | 5 | DESERT 2 |
| pEA1290_1114_1116_LTP | 100 | 69 | 21 | DESERT 2 |
| pEA1291_1117_1119_VRM | 100 | 55 | 70 | DESERT 2 |
| pEA1292_1120_1122_AII | 86 | 33 | 117 | DESERT 2 |
| pEA1293_1123_1125_KKS | 62 | 27 | 93 | DESERT 2 |
| pEA1294_1126_1128_GNN | 51 | 17 | 97 | DESERT 2 |
| pEA1295_1129_1131_RCW | 100 | 100 | 8 | DESERT 2 |
| pEA1296_1132_1134_RGC | 100 | 100 | 6 | DESERT 2 |
| pEA1297_1135_1137_GEI | 86 | 19 | 127 | DESERT 2 |
| pEA1298_1138_1140_GTL | 100 | 49 | 82 | DESERT 2 |
| pEA1299_1141_1143_LHC | 100 | 100 | 5 | DESERT 2 |
| pEA1300_1144_1146_WWD | 100 | 89 | 19 | DESERT 2 |
| pEA1301_1147_1149_CKL | 100 | 100 | 11 | DESERT 2 |
| pEA1302_1150_1152_VQP | 100 | 26 | 46 | DESERT 2 |
| pEA1303_1153_1155_LWK | 100 | 100 | 11 | DESERT 2 |
| pEA1304_1156_1158_SVW | 100 | 44 | 24 | DESERT 2 |
| pEA1305_1159_1161_RFL | 86 | 29 | 87 | DESERT 2 |
| pEA1306_1162_1164_RDL | 62 | 45 | 100 | DESERT 2 |
| pEA1307_1165_1167_ELE | 60 | 16 | 79 | DESERT 2 |
| pEA1308_1168_1170_IPF | 100 | 27 | 50 | DESERT 2 |
| pEA1309_1171_1173_DPA | 100 | 67 | 70 | DESERT 2 |
| pEA1310_1174_1176_IPL | 100 | 61 | 21 | DESERT 2 |
| pEA1311_1177_1179_LGI | 100 | 71 | 4 | DESERT 2 |
| pEA1312_1180_1182_YPN | 86 | 17 | 87 | DESERT 2 |
| pEA1313_1183_1185_EYK | 34 | 10 | 109 | DESERT 2 |
| pEA1314_1186_1188_SCC | 15 | 11 | 97 | DESERT 2 |
| pEA1315_1189_1191_YKD | 51 | 13 | 94 | DESERT 2 |
| pEA1316_1192_1194_TCT | 100 | 18 | 64 | DESERT 2 |
| pEA1317_1195_1197_RMF | 100 | 18 | 19 | DESERT 2 |
| pEA1318_1198_1200_IAA | 100 | 86 | 102 | DESERT 2 |
| pEA1319_1201_1203_LFT | 62 | 26 | 82 | DESERT 2 |
| pEA1320_1204_1206_IAK | 100 | 52 | 68 | DESERT 2 |
| pEA1321_1207_1209_TWN | 100 | 100 | 13 | DESERT 2 |
| pEA1322_1210_1212_QPK | 86 | 20 | 86 | DESERT 2 |
| pEA1323_1213_1215_CPT | 100 | 73 | 63 | DESERT 2 |
| pEA1324_1216_1218_MID | 51 | 16 | 91 | DESERT 2 |
| pEA1325_1219_1221_WIK | 100 | 89 | 25 | DESERT 2 |
| pEA1326_1222_1224_KMW | 100 | 21 | 8 | DESERT 2 |
| pEA1327_1225_1227_HIY | 62 | 22 | 24 | DESERT 2 |
| pEA1328_1228_1230_TME | 100 | 51 | 5 | DESERT 2 |
| pEA1329_1231_1233_YYA | 100 | 23 | 23 | DESERT 2 |
| pEA1330_1234_1236_AIK | 73 | 22 | 51 | DESERT 2 |
| pEA1331_1237_1239_NDE | 46 | 17 | 61 | DESERT 2 |
| pEA1332_1240_1242_FIS | 19 | 15 | 60 | DESERT 2 |
| pEA1333_1243_1245_FVG | 43 | 37 | 28 | DESERT 2 |
| pEA1334_1246_1248_TWM | 100 | 100 | 16 | DESERT 2 |
| pEA1335_1249_1251_KLE | 100 | 22 | 21 | DESERT 2 |
| pEA1336_1252_1254_TII | 62 | 19 | 9 | DESERT 2 |
| pEA1337_1255_1257_LSK | 100 | 14 | 56 | DESERT 2 |
| pEA1338_1258_1260_LSQ | 73 | 12 | 82 | DESERT 2 |
| pEA1339_1261_1263_EQK | 64 | 15 | 104 | DESERT 2 |
| pEA1340_1264_1266_TKH | 42 | 20 | 102 | DESERT 2 |
| pEA1341_1267_1269_RIF | 46 | 20 | 49 | DESERT 2 |
| pEA1342_1270_1272_SLI | 51 | 35 | 81 | DESERT 2 |
| pEA1343_1273_1275_GGN | 51 | 40 | 58 | DESERT 2 |
